# Myo1e/f regulate phagocytic podosomes to promote efficient cup closure in macrophages

**DOI:** 10.64898/2026.04.30.721640

**Authors:** Tyler C Paul, Yoseph M Loyd, Sharon E Chase, Thomas W O’Connor, Chad M Hobson, Rachel M Lee, Daan Vorselen, Mira Krendel

## Abstract

Phagocytosis requires coordinated remodeling of the actin cytoskeleton to drive target engulfment. Class I myosins Myo1e and Myo1f contribute to Fc-receptor-mediated phagocytosis. Using CRISPR-edited RAW 264.7 macrophages, we show that cells lacking Myo1e/f exhibit reduced uptake of IgG-coated beads. Lattice-light-sheet and confocal imaging revealed distinct F-actin architectures corresponding to the various stages of cup progression, including basal podosome-like adhesions, individual actin nodes along the rim of the cup, and a highly contractile, actin-rich phagocytic ring. In Myo1e/f-deficient cells, podosome formation was diminished, and the phagocytic ring formed prematurely, which was often accompanied by the strong deformation of the phagocytic targets and repeated engulfment attempts. Myo1e/f localized both to podosome-like structures and to the inner surface of the phagocytic ring, non-muscle myosin II (NM2) localized to the outer surface, and the absence of Myo1e/f correlated with the broader distribution of NM2. In addition, Myo1e/f-deficient macrophages exhibited increased trogocytosis of antibody-opsonized HL-60 cells. These results suggest that Myo1e/f coordinate spatial and temporal transitions between protrusive and contractile actin networks, thereby ensuring efficient phagocytic cup progression. Our findings highlight a dual role for Myo1e/f in adhesion regulation and force balance during macrophage phagocytosis.

## Introduction

Phagocytosis is a complex and dynamic process in which specialized immune cells, termed professional phagocytes, engulf pathogens, apoptotic cells, and cellular debris (Boada-Romero et al., 2020; Depierre et al., 2025; Flannagan et al., 2012). This process is initiated when receptors on the phagocyte bind to specific ligands on potential phagocytic targets, including opsonins (surface-marking proteins) such as complement proteins or antibodies (Greenberg and Grinstein, 2002).

In Fc-receptor (FcR) mediated phagocytosis, the Fc portion of an antibody bound to a target engages FcRs on the phagocyte surface (Daëron, 1997). Following this initial contact, FcRs cluster to form adhesions with the target (Sobota et al., 2005). Subsequently, the cell membrane extends around the target, forming the actin-rich phagocytic cup, which expands until the target is fully internalized into a phagosome (Greenberg and Grinstein, 2002). The contents of the phagosome are then bound for degradation through fusion with lysosomes (Gordon, 2016).

Recent studies have identified podosome-like structures (dynamic actin-based adhesions) during the formation and progression of the phagocytic cup (Barger et al., 2019; Ostrowski et al., 2019; Vorselen et al., 2021). Podosomes consist of a core composed of branched actin filaments (F-actin) surrounded by a ring composed of adhesion proteins such as paxillin, vinculin, and talin and are involved in cell adhesion and migration (Linder et al., 2023). They are characteristic of hematopoietic cells such as macrophages (Linder et al., 1999) and osteoclasts (Destaing et al., 2003). Structures that are similar in size, protein composition, and protrusive activity to podosomes have been detected at multiple stages of phagocytosis, from initial target adhesion (Barger et al., 2019) to cup extension and closure (Ostrowski et al., 2019; Vorselen et al., 2021), where they may contribute to protrusive force generation within the cup.

Class-I myosins, single-headed myosin motor proteins, are commonly found in association with branched F-actin. Using a positively charged tail homology 1 (TH1) domain, they bind to anionic phospholipids such as PI(4,5)P_2_ in the plasma membrane (Adams and Pollard, 1989; Feeser et al., 2010; Hayden et al., 1990). By interacting with branched actin networks via their motor domains, they generate a connection between actin filaments and the plasma membrane. Two members of the class I myosin family, myosin 1e and myosin 1f (Myo1e/f), localize to actin-rich, podosome-like protrusive structures in phagocytic cups (Barger et al., 2019) and to the base of conventional podosomes (Cervero et al., 2025). Previous studies have shown that Myo1e/f help maintain membrane tension (Barger et al., 2019; Nambiar et al., 2009), regulate F-actin organization during phagocytosis, and promote efficient target engulfment (Barger et al., 2019; Barger et al., 2022). Our previous study using RAW 264.7 macrophages highlighted the following observations: Myo1e/f localize to the leading edge and inner surface of the phagocytic cup and are concentrated at the membrane-attached tips of actin nodes (“teeth”) (Barger et al., 2019). Complementary experiments in the same study, using bone marrow derived macrophages (BMDM) from Myo1e/f dKO mice, have allowed us to identify a defect in phagocytic cup closure (delayed cup closure) in the absence of Myo1e/f; in a separate study we have observed that Arp2/3-nucleated actin nodes and non-muscle myosin-II generate complementary forces (protrusive and constrictive, respectively) during phagocytosis. However, since BMDMs exhibit a relatively gentle force signature and fairly uniform actin distribution, with less pronounced actin structures at the rim of the cup (Barger et al., 2022), using these primary cells made it difficult to resolve how the loss of Myo1e/f affects specific actin structures within phagocytic cups, such as protrusive actin teeth and contractile actin rings enriched in myosin II. Wild-type RAW 264.7 macrophages have better-defined actin “teeth” and produce stronger protrusive forces than wild-type mouse BMDMs. Therefore, in order to follow up on our observations in the Myo1e/f dKO BMDMs and precisely define the spatiotemporal contributions of Myo1e/f to phagocytic cup architecture, here we employed the RAW 264.7 macrophage model, which provides a highly reproducible and distinct pattern of target deformation and actin dynamics ideal for mechanical studies.

To precisely dissect spatial and temporal transitions between the different stages of phagocytic uptake, we used CRISPR/Cas9 to generate Myo1e/f double-knockout RAW 264.7 macrophages. We then investigated how the loss of Myo1e/f alters actin reorganization during phagocytosis of deformable acrylamide-coacrylic acid microparticles (DAAMPs), utilizing live-cell lattice light sheet microscopy (LLSM) and fixed-cell confocal imaging. We find that dKO of Myo1e/f changes the balance between formation of individual protrusive podosome-like structures and their assembly into contractile rings within phagocytic cups, likely leading to reduced target uptake. These findings define a mechanistic role for Myo1e/f in orchestrating the cytoskeletal transitions that enable efficient phagocytic cup progression.

## Results

### Generation and characterization of the Myo1e/f double knockout RAW2C4.7 cells

To investigate the role of Myo1e/f in phagocytosis, we generated Myo1e/f dKO RAW264.7 immortalized murine macrophages using CRISPR/Cas9 gene editing performed by Synthego. The editing introduced frameshift mutations in exon 2 of both *Myo1e* and *Myo1f,* leading to premature stop codons (Fig.1A). Following limiting-dilution cloning, two independent clones (B1 and D9) carrying stop codons in both alleles of *Myo1e* and *Myo1f* genes were identified and returned to our laboratory. The B1 clone was used as the main dKO cell line in this study.

**Figure 1.**
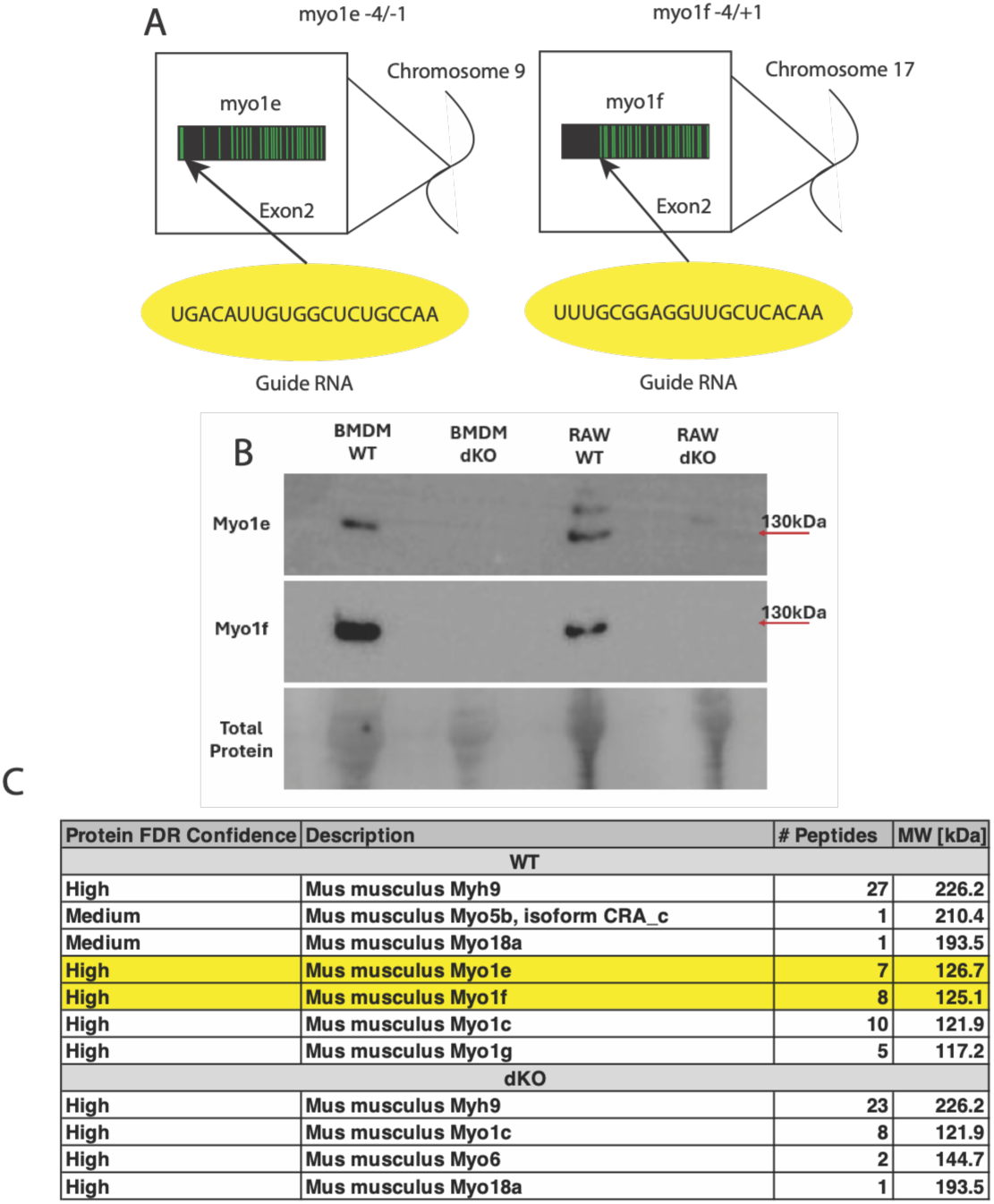
CRISPR-Cas3 targeting of exon 2 in Myo1e and Myo1f generates myosin double knockout in RAW2C4.7 macrophages. **(A)** Schematic of CRISPR–CasS editing strategy. Guide RNAs targeting exon 2 of Myo1e (left) and Myo1f (right) (yellow) introduced small indels (−4/−1 for Myo1e and −4/+1 for Myo1f), resulting in frameshifts and premature stop codons in both genes. The indels shown correspond to the edits identified in the RAW dKO clone (B1) that was used for most experiments in this study. **(B)** Western blot validation of Myo1e/f loss. Comparison of the edited clone (B1) with WT RAW 2C4.7 cells and bone marrow–derived macrophages (BMDMs) confirmed loss of Myo1e and Myo1f expression. Arrows indicate the expected molecular weights of Myo1e and Myo1f. There are additional bands unique to RAW cells that are recognized by the anti-Myo1e antibody. SYPRO Ruby total protein stain was used as a loading control. **(C)** Mass spectrometry confirmation of Myo1e/f knockout. SDS–PAGE gel regions corresponding to 50–250 kDa were excised and analyzed. Myo1e and Myo1f peptides were absent in dKO samples but detected in WT controls. False discovery rate (FDR) confidence, peptide counts, and predicted molecular weights are shown.

Western blotting with antibodies against Myo1e or Myo1f confirmed the absence of Myo1e and Myo1f protein bands in dKO cells compared to wild-type (WT) controls (Fig.1B, S1A). In addition to the expected Myo1e band observed only in WT lysates, an additional band recognized by the anti-Myo1e antibody was detected in both WT and dKO cells. To determine whether this band represented Myo1e or an alternatively spliced Myo1e isoform, we excised the 50-250 kDa region from SDS-PAGE gels and performed proteomic analysis. Myo1e and Myo1f peptides were detected in WT but not dKO samples, confirming the specificity of the knockout (Fig.1C).

To further control for potential clonal variability, we performed a second round of limiting-dilution cloning from the edited RAW 264.7 pool. Twenty-two additional clones were isolated, and eleven were screened by Western blotting to identify additional Myo1e/f-deficient lines (Fig. S1D).

### Myo1e/f deficiency results in decreased uptake of IgG-coated beads

To determine how Myo1e/f loss affects phagocytosis, we performed a phagocytic efficiency assay using WT and Myo1e/f dKO (clone B1) RAW 264.7 cells. Cells were incubated with 9 μm IgG-coated polystyrene beads that were centrifuged onto the cell surface to synchronize uptake. Phagocytosis was stopped by fixation at specified time points, and the exposed surface of the beads was detected by staining with fluorescent secondary antibodies to determine what fraction of all beads was engulfed at each time point (Fig. 2A,B). Wide-field fluorescence images were used to measure the portion of the bead surface that was covered by the cell and therefore inaccessible to secondary antibody staining as described in the Materials and Methods. This analysis revealed a bimodal distribution of beads, with the majority of beads either mostly exposed to the outside or mostly protected from secondary antibody staining (Fig. S1B). Beads that were more than 85% protected from the secondary antibody staining were considered fully engulfed and used to calculate phagocytic uptake efficiency (fraction of all beads in each field of view that was fully engulfed). A pronounced reduction in phagocytic efficiency was observed at the 15 and 30-minute timepoints in the Myo1e/f dKO cells compared to WT controls (Fig. 2C,D, S1C). To confirm that the phagocytic defect was not limited to a single dKO clone, we performed the same assay with several additional Myo1e/f-deficient clones generated from the edited cell pool. Each of these clones exhibited some degree of the phagocytosis defect (Fig.S1C,E).

**Figure 2.**
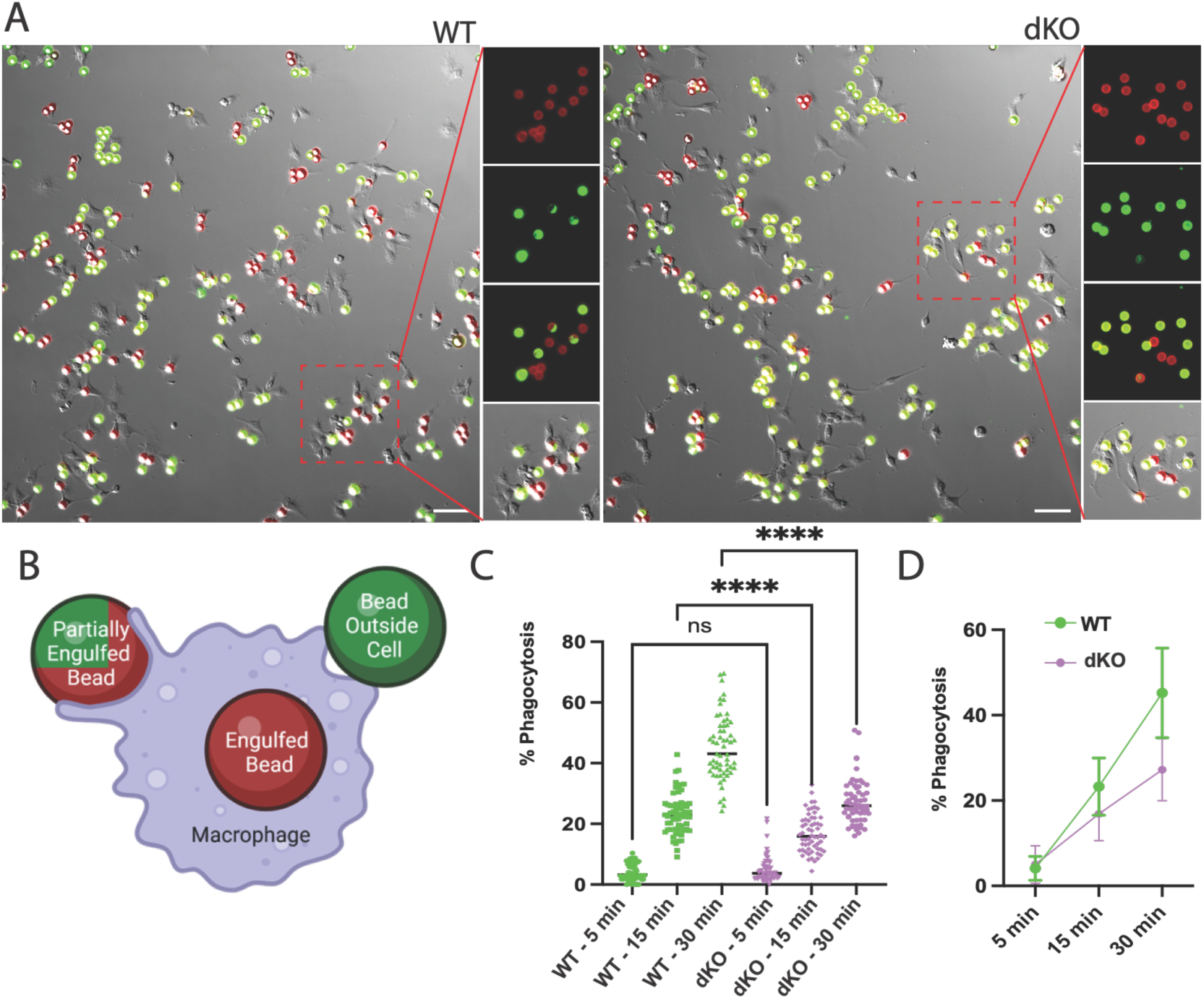
Myo1e/f double knockout impairs phagocytic efficiency in RAW 2C4.7 macrophages. (**A**) Representative images from the 30-min timepoint in the phagocytic efficiency assay using polystyrene beads (9 μm diameter). Merged fluorescence and DIC images show an increased number of fully engulfed beads (red) in WT RAW 2C4.7 macrophages (left) compared to Myo1e/f dKO cells (right, clone B1). A 100 × 100 μm region (red box) highlights a representative area with individual analysis channels: all beads (red), surface-exposed/unengulfed beads (green), and merged views (2-channel to highlight partial engulfment; 3-channel to show total cells vs phagocytosing cells). Scale bar: 50 μm. (**B**) Cartoon illustrating the process of identification of fully engulfed vs. partially enclosed beads. In merged images, fully engulfed beads appear red, fully unengulfed beads appear green, and partially engulfed beads show combined red/green signal. (**C**) Analysis of phagocytic efficiency in WT vs. dKO cells. Phagocytic efficiency (ratio of beads that were more than 85% enclosed by the cells to all beads) was quantified for each field of view as described in Materials and Methods and expressed as %. Each data point represents one field of view. N= 3 independent experiments. One-way ANOVA test shows a significant decrease in phagocytic efficiency in dKO cells at 15 and 30 mins compared with WT cells. (**D**) Same data as in **C** are displayed as mean values for all fields of view for each time point. Error bars indicate SD. Statistical significance: ns: p = 0.S7S5, ****: p < 0.0001. n =54 fields of view for dKO at 5 min, n =58 fields of view for WT at 5 min, n =59 fields of view for WT at 30 min, n =60 fields of view for each of the other conditions, N =3 for all conditions. The number of beads analyzed for each condition was over 5500.

To test whether the reintroduction of Myo1e or Myo1f could rescue the phagocytic defects observed in dKO cells, we transfected dKO cells with Halo-tagged Myo1e or Myo1f constructs (Fig. S1F,G) and performed the phagocytic efficiency assay at the 30-minute timepoint. Previously we have demonstrated that single KO of Myo1e or Myo1f does not produce a significant phagocytosis defect, indicating that the two myosins may play redundant roles in phagocytosis (Barger et al., 2019). Re-expression of either protein only partially restored phagocytic efficiency in dKO cells (Fig. S1H); the incomplete rescue could be explained, at least in part, by the relatively low transfection efficiency for myosin constructs (Fig. S1F,G). On the other hand, expression of Halo-tagged Myo1e or Myo1f reduced phagocytic efficiency in the WT cells (Fig. S1H), which may indicate that myosin overexpression disrupts existing protein-protein interactions and interferes with the productive actin-membrane remodeling driven by the endogenous myosins.

### Dynamic F-Actin Transitions During Phagocytosis

We next examined actin reorganization during phagocytic cup progression and the corresponding changes in the deformation of phagocytic targets using two complementary approaches: live-cell imaging and imaging of fixed phagocytic cups. First, we used lattice light-sheet microscopy (LLSM) to observe phagocytosis in live macrophages. Fluorescently labeled deformable acrylamide-co-acrylic acid microparticles (DAAMPs) served as phagocytic targets, allowing simultaneous observation of actin dynamics and target deformation by LLSM (Mali et al., 2026; Vorselen et al., 2020). Using LLSM, we imaged WT RAW264.7 macrophages expressing mEmerald–Lifeact to label F-actin as they engulfed Alexa-647– cadaverine labeled DAAMPs opsonized with IgG (Fig.3A-D, Fig.S2A-C, Supp Videos 1,3). In the initial phase of phagocytosis, broad actin-rich ruffles and lamellipodia were formed during particle engagement and cup formation. As uptake progressed, discrete actin-rich foci became resolvable within or adjacent to this continuous cup-associated actin network. A subset of these foci coincided spatially and temporally with localized deformation of the particle surface (Fig.3 and S2, white arrows). Because individual foci could not always be distinguished against the intense actin signal within the cup, arrows indicate only the most clearly resolved examples. Some of the F-actin-rich hotspots coinciding with regions of deformation were located at the base of the beads. We refer to these early structures as basal podosomes. As the cup advanced over the bead, additional F-actin hotspots appeared near the leading edge (Fig. 3B,C,D, white arrows). These structures varied in the extent of bead deformation but consistently coincided with localized indentations. At later stages of engulfment, a dense band of F-actin, referred to here as the phagocytic ring, formed at the membrane’s leading edge and coincided with intense deformation of the bead surface (Fig.3, Fig.S2B,C, blue arrows). The phagocytic ring persisted through target enclosure, after which F-actin reorganized into a cap-like structure over the top of the bead (Fig.3A-D, Fig.S2B,C, orange arrows), producing marked deformations likely associated with pushing the newly internalized phagosome toward the cell interior. This bright actin cap is reminiscent of the burst of actin assembly previously observed following phagosome closure in complement receptor-mediated phagocytosis, which was also proposed to propel the phagosome inward (Jaumouillé et al., 2019).

**Figure 3.**
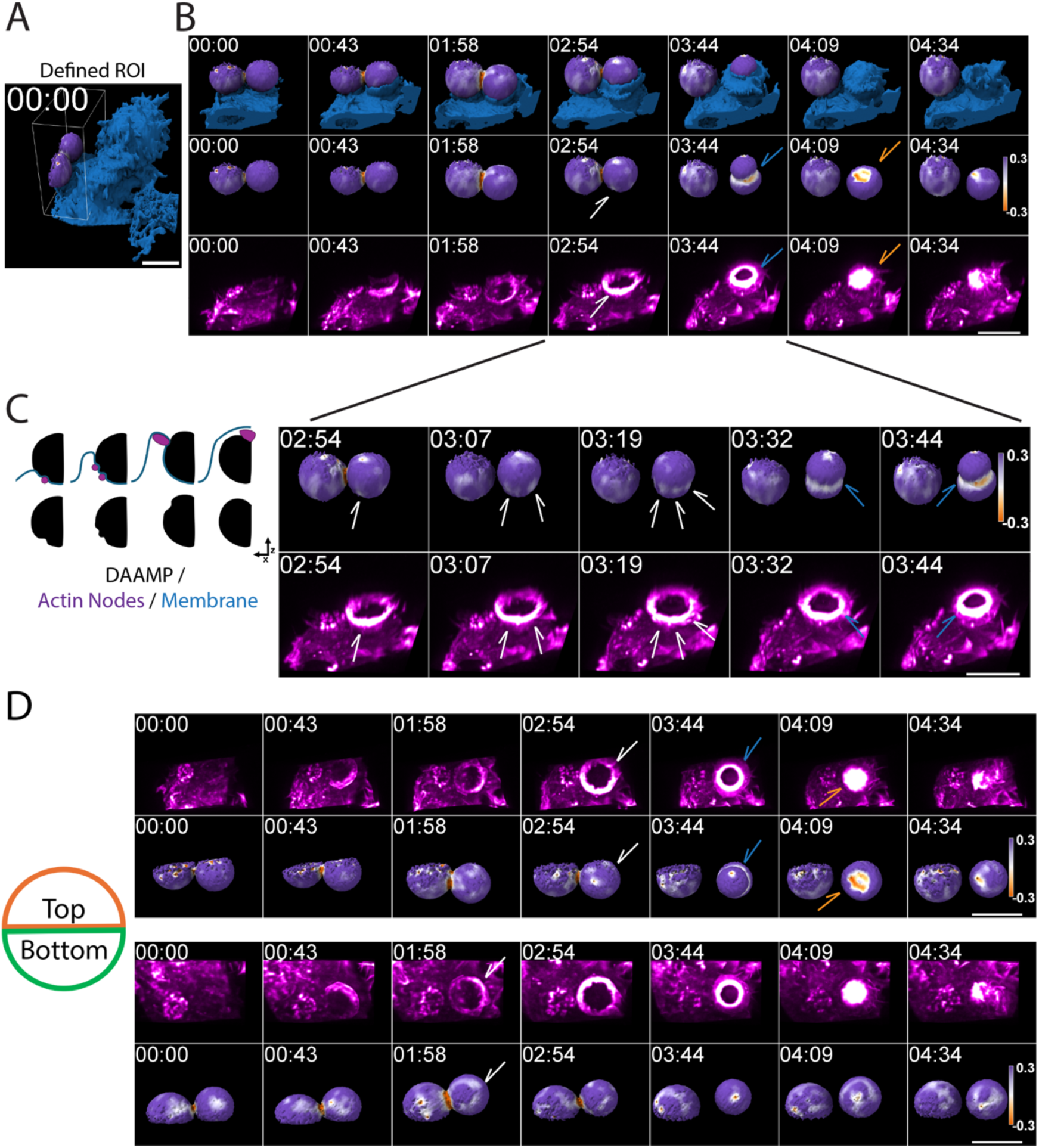
Typical progression of F-actin reorganization and bead deformation during WT phagocytosis visualized using live-cell LLSM. **(A)** Overview of macrophages expressing mEmerald–Lifeact (blue isosurface) interacting with deformable IgG-coated DAAMPs, represented as an isosurface with a diverging purple-orange colormap representing local curvature measurements between -0.3 and 0.3 steradians, with purple representing convex regions of the bead and orange indicating indented regions of the bead. Wireframe box indicates the region of interest (ROI) corresponding to the montage shown in **(B, C,D)**.**(B)** The time-lapse montage depicts the right bead undergoing complete engulfment and the left bead remaining at the early attachment stage. Cell shape is represented by the blue isosurface derived from mEmerald-Lifeact labeling, DAAMPs are shown using the purple-orange curvature color map as described for panel **A**, and F-actin is visualized using a volumetric maximum intensity projection (MIP) paletted as magenta-white heatmap to highlight areas of actin enrichment. **(C)** Left panel: Schematic illustrating bead (black) deformations driven by F-actin-rich podosome-like structures (purple) and macrophage membrane (blue) dynamics during antibody mediated phagocytosis. Right panel: A portion of the time-lapse movie shown in **B** is displayed at higher magnification and with shorter time intervals. **(D)** Same sequence as in **B** is shown in two additional viewing orientations, with the view from the top or the bottom, as indicated. In all panels white arrows point to actin hot spots that coincide with distinct spot-like deformations of the bead surface, blue arrows highlight phagocytic rings that colocalize with continuous belt-like areas of deformation, and orange arrows highlight cap-like actin spots that form upon cup closure. Scale bars, 10 μm.

To verify our observations, we used confocal imaging of fixed cells to collect a more extensive dataset of actin organization and target deformations at different stages of cup closure (Fig.4). Using confocal imaging of WT cells fixed during engulfment of DAAMPs, we examined the appearance of individual actin structures, labeled with phalloidin, at different stages of phagocytosis (Fig.4A). In phagocytic cups corresponding to the stages from the initial bead contact to approximately 20% of bead engulfment (as determined using the secondary antibody staining of the exposed bead surface), we observed distinct basal F-actin nodes (presumably phagocytic podosomes) that coincided with the sites of bead surface indentation (Fig.4B, green arrows). At approximately 20-30% cup progression, we observed the presence of bright actin nodes located near the leading edge of the cup, consistent with the phagocytic actin “teeth” described previously (Vorselen et al., 2021) (Fig.4C). Finally, we observed continuous phagocytic actin rings in many cells that had engulfed 50% or more of the bead (Fig.4D). While the phagocytic rings seemed to be most often present at these stages, there were rare instances where we found rings at earlier stages, with one example found at 12% engulfment. Observed phagocytic rings were often proximal to continuous rings of bead deformation (Fig.4D). Bead sphericity measurements showed a decrease, indicating a period of heavy deformation, at stages correlating with the appearance of actin teeth (20-30%) and the phagocytic ring (∼50%) (Fig.4E).

Together, our observations from the LLSM and confocal imaging of WT RAW264.7 macrophages support a model for phagocytic cup progression where basal podosomes establish initial target contacts, phagocytic actin teeth maintain attachment as the cup advances, and the phagocytic ring emerges around mid-engulfment (∼50%), persisting through closure (Fig.4F).

**Figure 4.**
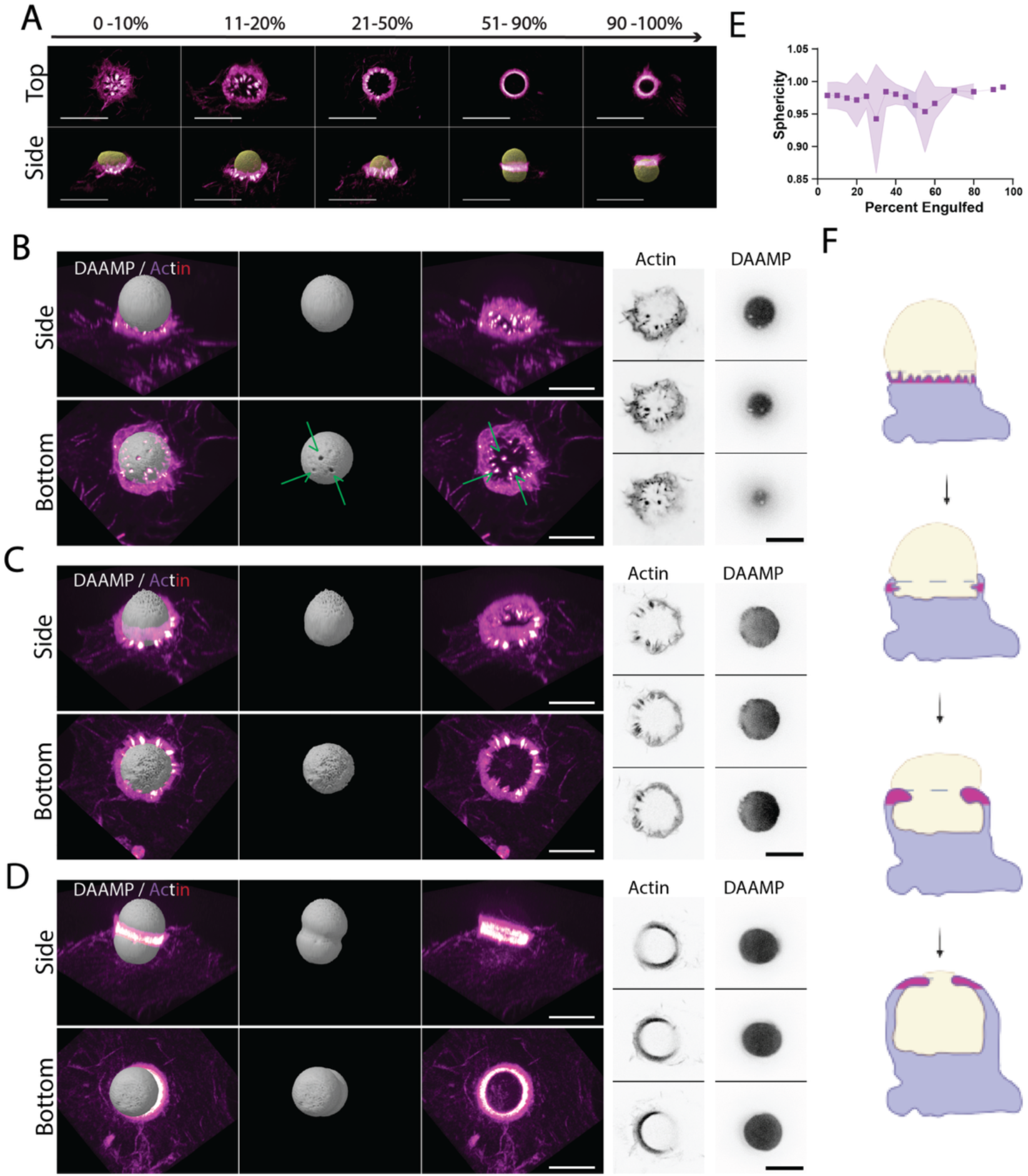
Fixed-cell imaging reveals three distinct types of actin structures during phagocytosis of DAAMPs by WT RAW 264.7 cells. **(A)** Representative phagocytic cups at various stages of engulfment. Top-down (top) and side-view (bottom) images of fixed RAW 2C4.7 WT macrophages stained with phalloidin (magenta heatmap) show the characteristic organization of F-actin around IgG-coated DAAMPs (yellow) from early attachment (left) to late-stage enclosure (right). Scale bars: 10 μm. **(B–D)** Montages illustrating basal podosomes (B, green arrows), actin teeth (C), and the phagocytic ring (D). For each structure, side and bottom-up views show F-actin (magenta heatmap) relative to the bead (gray). Grayscale panels on the right show representative z-slices from the F-actin and bead channels. Scale bars: 5 μm. **(E)** Bead sphericity changes during phagocytic progression. Sphericity measurements, binned in 5% increments of phagocytic progression (engulfment), show characteristic dips at ∼25–30% and ∼55–C0%, corresponding to bead deformation associated with the actin teeth and the phagocytic ring, respectively (purple dots indicate mean values, shaded areas indicate SD, n = 160 beads). **(F)** Schematic depicting actin reorganization during bead uptake. The diagram highlights the spatial organization of basal podosomes, actin teeth, and the phagocytic ring (magenta) relative to the macrophage (lavender) and bead (beige) throughout phagocytic progression.

### Absence of Myo1e/f causes premature formation of the phagocytic actin ring and stalled cup closure

After characterizing the typical progression of phagocytosis and F-actin organization in WT macrophages, we next examined Myo1e/f dKO cells to identify alterations that might underlie their reduced phagocytic efficiency. Under identical LLSM conditions, some of the dKO macrophages exhibited a distinctive behavior not often observed in WT cells: they formed the phagocytic ring prematurely at an early stage of cup closure, after which the cup receded, and the cell reattempted engulfment (representative sequences shown in Fig. 5A-D, Fig.S3, Supp. Videos 2,4). Quantification of the number of time-lapse sequences demonstrating repeated formation and contraction of phagocytic cups in the WT and dKO cells produced the following distribution: of 11 complete engulfment sequences in the WT cells, only 1 sequence demonstrated repeated cup formation and multiple engulfment attempts; of 13 complete engulfment sequences in the dKO cells, 3 demonstrated early contractile ring formation and repeated engulfment attempts. Due to the challenges of live-cell observations, the number of completed engulfment attempts was limited and did not allow for sufficient data collection for statistical analysis. Therefore, we performed analysis of fixed cells to determine whether specific types of phagocytic actin structures were observed with different frequency in WT and dKO macrophages.

In order to determine the approximate distribution of actin structures at various stages of phagocytic cup closure, fixed phagocytic cups were binned based on the extent of target engulfment. From observations of 197 WT and 192 dKO RAW 264.7 phagocytic cups, the phagocytic ring appeared more frequently at early stages of phagocytosis in dKO cells (Fig. 6A-B). The phagocytic ring was present in 25% of dKO cups prior to 50% engulfment compared to only 11% in WT cups. Using the DAAMParticle_Shape_Analysis Matlab functions we analyzed bead sphericity in 160 WT and 151 dKO phagocytic cups and observed a reduction in sphericity in dKO cells prior to 50% closure (Fig. 6C-D). This decrease in sphericity likely reflects premature formation of the phagocytic ring and the associated dramatic bead deformation.

Fixed-cell analysis also revealed several additional deviations from the WT phenotype. dKO cells formed distinct basal podosomes at a lower frequency than WT cells (Fig. 6E): 60% of WT cups displayed distinct basal podosomes compared to 32% of dKO cups. Interestingly, some deformation was still detectable at the bead base even in the absence of clear podosome-like structures. “Actin teeth” (individual podosome-like structures located along the cup rim) were rarely present in dKO cups (Fig. 6F), appearing in only 6% of dKO cups at any stage compared to 35% of WT cups. Consistent with the reduced formation of phagocytic podosomes, a greater proportion of dKO cells displayed a loose association with their targets, defined by the observable separation between the actin lamellipodia at the rim of the phagocytic cup and the bead surface (Fig. 6G, Fig.S5A-J). Although this feature was occasionally observed in WT cells, it was much more frequent in dKO cells (63% of dKO cups at any stage compared to 18% of WT cups).

**Figure 5.**
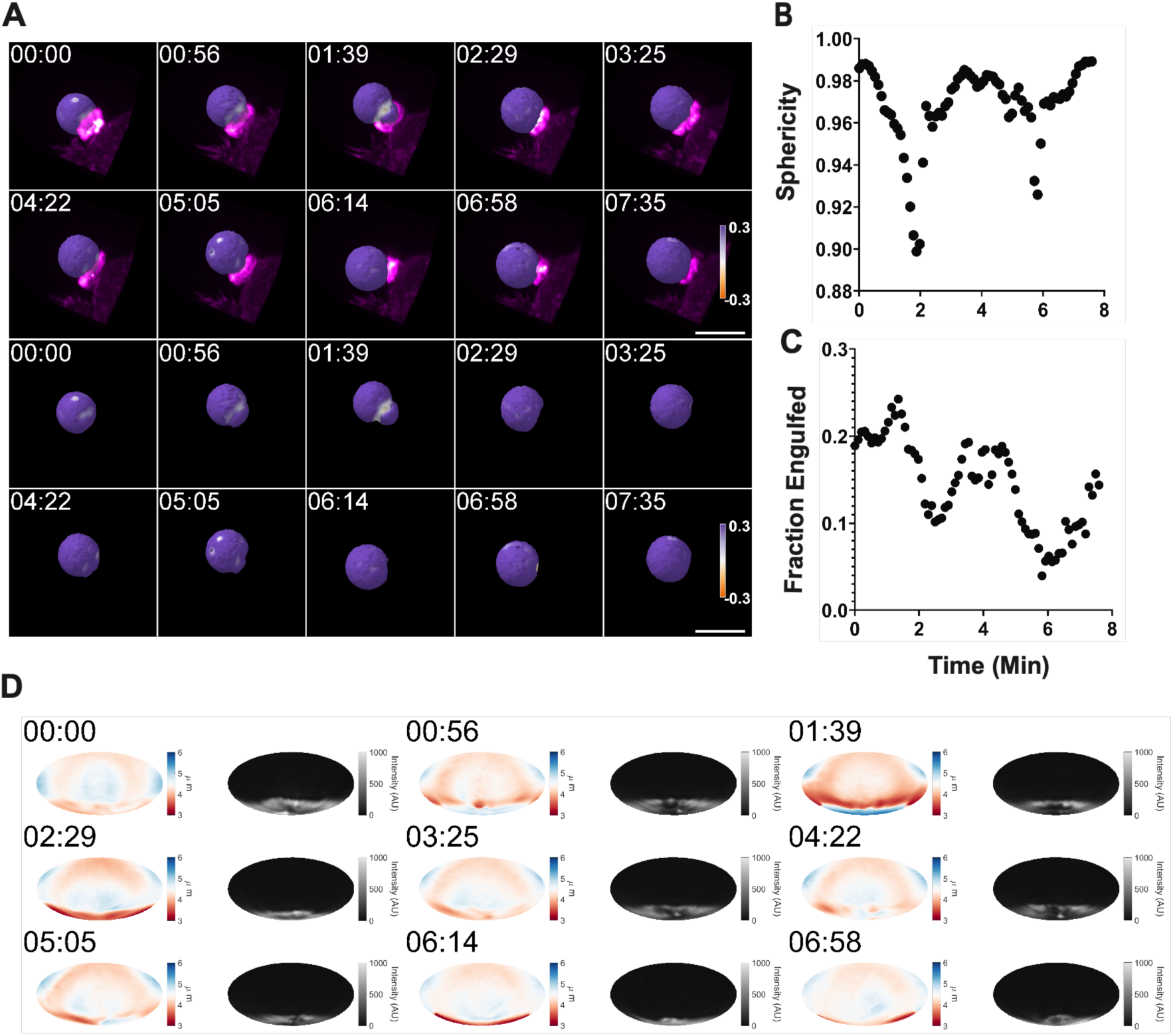
Loss of Myo1e/f leads to premature contraction and stalling of the phagocytic cup. **(A)** Time series of a Myo1e/f dKO macrophage expressing mEmerald–Lifeact (F-actin; magenta intensity map) unsuccessfully attempting to engulf an IgG-coated DAAMP. The DAAMP surface is rendered with a purple-to-orange diverging colormap encoding vertex-based angular curvature derived from the triangulated surface. Lower panels show the bead alone to highlight surface deformation. In this image sequence, the macrophage attempts several times to engulf the DAAMP, while indenting the particle. Scale bar: 10 μm. **(B)** A graph of sphericity of the bead over time for the event shown in panel A highlights repeated constriction of the cup followed by relaxation. **(C)** Fractional bead engulfment over time for the same event. A drop in bead sphericity precedes the reduction in engulfment fraction, indicating premature contraction and subsequent cup stalling or cup retraction. **(D)** Mollweide projections showing bead surface distance to centroid ranging between 3 µm and 6 µm (with red illustrating areas of bead indentation and blue indicating bead surface areas that are not compressed) and corresponding F-actin distribution (right) at each timepoint (excluding the final frame).

**Figure 6.**
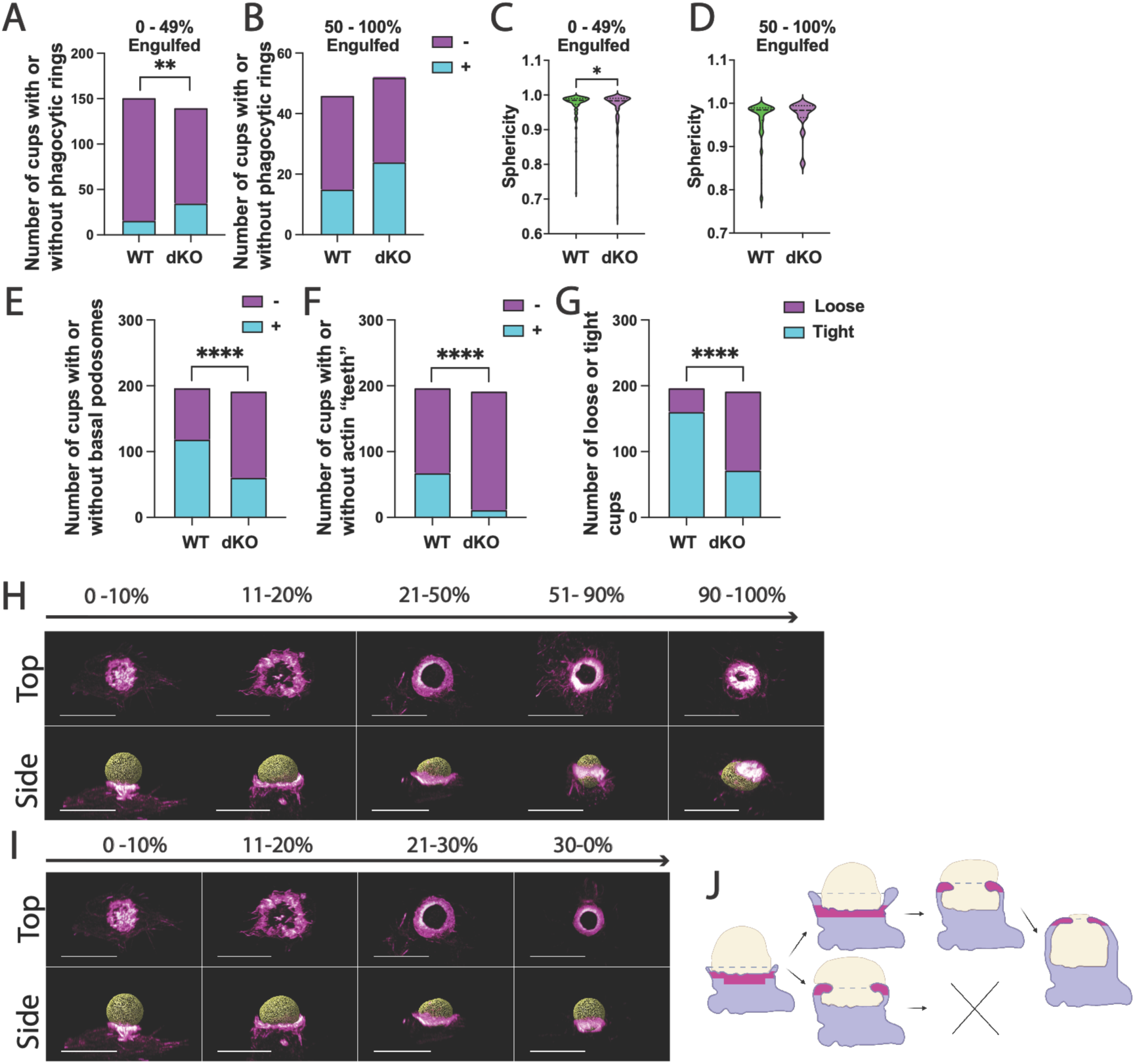
Myo1e/f dKO disrupts the spatiotemporal regulation of F-actin structures during phagocytosis, contributing to delayed or failed particle uptake. (**A–B**) Frequency of phagocytic ring formation before (A) and after (B) 50% cup closure. Parts-of-a-whole bar graphs show the number of fixed RAW 264.7 WT and Myo1e/f dKO macrophages with (cyan) or without (magenta) a phagocytic ring. Replicates WT n = 151 in A, 46 in B, N = 4; dKO n = 140 A, 52 in B, N = 4. ** p= 0.0018 (Fisher’s exact test, A). No statistically significant difference was detected in B. (**C–D**) Bead sphericity in fixed cells before (C) and after (D) 50% cup closure. Violin plots reveal a significant decrease in bead sphericity, indicating pronounced deformation, in dKO cups prior to reaching 50% progression. (C) WT n = 137, dKO n = 139; N=4; (D) WT n = 23, dKO n = 12; N=4. * p < 0.05 (unpaired t test). (**E–G**) Number of cups with and without basal podosomes (E), with and without actin teeth (F), and loose/tight cups (G). WT n = 197, N = 4; dKO n = 192, N = 4.**** p < 0.0001, Fisher’s exact test. (**H–I**) Representative fixed phagocytic cups consistent with the proposed successful and failed outcomes in dKO cells, with % indicating the extent of bead coverage by the cell. Top-down and side-view images of phalloidin-labeled F-actin (magenta heatmap) and Alexa-647–cadaverine–labeled DAAMPs (yellow) show: (**H**) Successful fate: phagocytic ring formation in late stages, followed by continued progression. (I) Failed fate: phagocytic ring formation in early stages, associated with subsequent stalling. Scale bars: 10 μm. (J) Schematic of phagocytic fates in Myo1e/f dKO cells. Diagram illustrates podosome structures (magenta) relative to the macrophage (lavender) and DAAMP (beige).

Integrating findings from both LLSM and fixed-cell imaging, we propose a model of phagocytosis in dKO macrophages that involves two distinct fates (Fig. 6H-J). In one outcome, the phagocytic ring forms at an appropriate stage, and the cup progresses to full closure (Fig. 6H). In the alternate fate, the phagocytic ring forms prematurely, and cup progression is stalled (Fig. 6I).

### Myo1e/f localize to the phagocytic ring and podosomes at the base of the phagocytic cup

**Figure 7.**
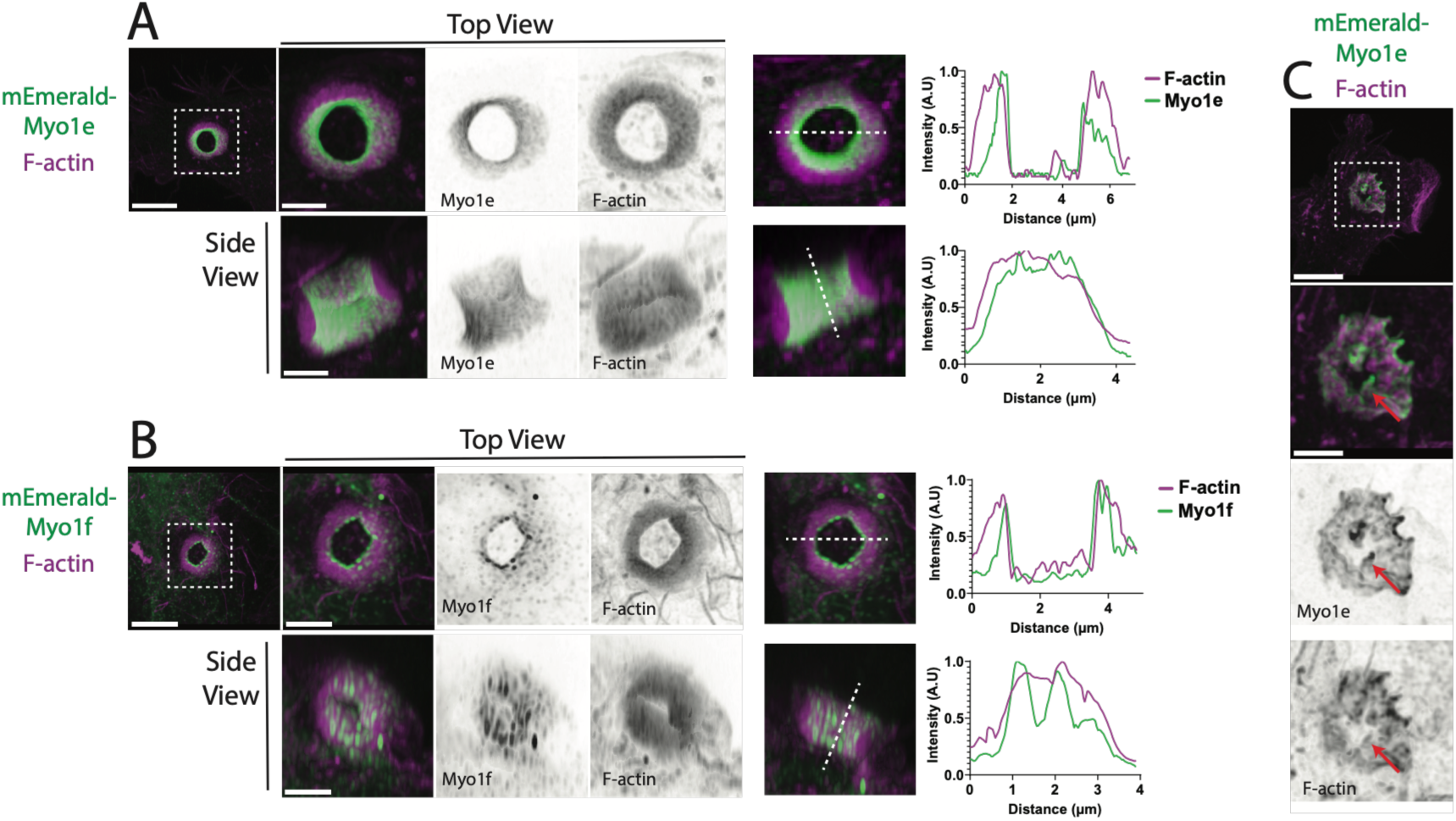
Myo1e and Myo1f localize to key podosome-like F-actin structures within the phagocytic cup. (**A-B)** Myo1e and Myo1f enrichment in the phagocytic ring. RAW2C4.7 WT cells expressing mEmerald-Myo1e (A) or mEmerald-Myo1f (B) (green) show enrichment along the inner surface of the phagocytic cup. F-actin was labeled with phalloidin (magenta). Shown are 3D projections with the bead excluded for clarity, displayed as top-down views (top) and side views (bottom). Line profiles corresponding to the dotted line are shown at right. Scale bars: main panels, 5 μm; zooms, 2 μm. **(C)** Myo1e localization to basal podosomes in early-stage cups. mEmerald-Myo1e (green) is enriched at the tips of F-actin puncta (magenta) within basal podosomes. Arrow indicates a basal podosome. Scale bars: main panel, 5 μm; zoom, 2 μm.

We next examined the localization of Myo1e and Myo1f within the phagocytic cup to gain insight into how their loss may lead to altered actin organization. Based on the previous reports of Myo1e association with adhesion and actin-rich structures, we hypothesized that both Myo1e and Myo1f would concentrate at the tips of podosomes at the base of the cup and along the leading edge of the cup when the phagocytic ring is present. To test this, we transfected WT and dKO macrophages with mEmerald-tagged Myo1e or Myo1f and performed phagocytosis assays, fixing cells 10 minutes after bead addition and staining F-actin with phalloidin.

Confocal imaging revealed that Myo1e and Myo1f exhibited overlapping localization patterns. Both proteins were distributed throughout the inner surface of the phagocytic ring at the membrane–bead interface (Figs. 7A-B) and enriched at the tips of podosomes at the base of the phagocytic cup (Fig. 7C, Fig.S4A). We confirmed previous reports that Myo1e/f localize to the tips of phagocytic actin teeth (Vorselen et al., 2021) (Fig.S4B-C). This localization pattern was consistent in both WT cells and in dKO cells reexpressing either construct, indicating that Myo1e and Myo1f redundantly associate with key actin structures that govern cup progression.

### NM2 localization becomes more diffuse in the absence of Myo1e/f

**Figure 8.**
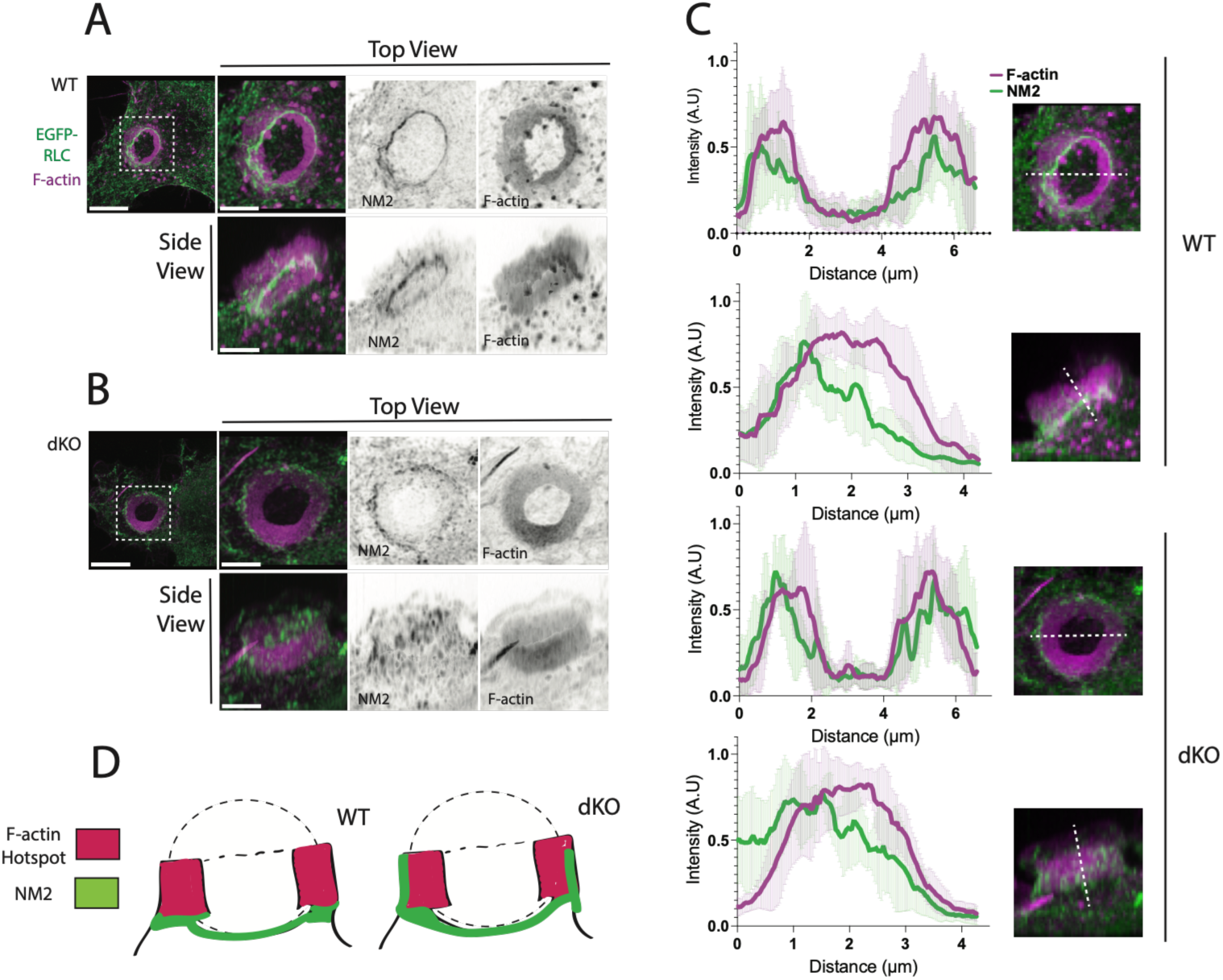
NM2 localization is altered in Myo1e/f dKO phagocytic rings. (A-B) NM2 localization relative to F-actin in the phagocytic actin ring in RAW2C4.7 WT and dKO macrophages. In RAW 2C4.7 WT macrophages expressing EGFP-Regulatory Light Chain (EGFP-RLC; green), non-muscle myosin II (NM2) forms a concentrated band positioned at the outer edge and base of the phagocytic actin ring (A). F-actin is labeled with phalloidin (magenta). In dKO cells (B), NM2 remains enriched at the outer ring but displays a more diffuse distribution along the vertical axis, rather than the sharply defined band observed in WT cells. Scale bars, 5 μm, zoom scale bars, 2 μm.**(C)** Line profiles of NM2 and F-actin distribution in the phagocytic actin ring. Top-down and side-view intensity profiles were collected from C phagocytic rings per cell line and plotted as normalized fluorescence intensity (y-axis) versus position (x-axis, μm), with error bars representing standard deviation from C measurements. In top-down views of both WT and dKO rings, NM2 (green) localizes to the outer edges of F-actin ring (magenta). In side-view profiles, WT NM2 signal peaks at the base of the ring and rapidly decreases towards the top. For dKO rings, NM2 localization more closely resembles that of F-actin, with a broad peak of fluorescence intensity. Shaded green or magenta areas indicate the standard deviation for NM2 and F-actin, respectively.**(D)** Schematic of NM2 localization in phagocytic actin rings in WT or dKO macrophages. The phagocytic ring is represented in magenta and NM2 is represented in green, illustrating the spatial restriction of NM2 to the outer-basal band in WT cells and the broader, more diffuse distribution in dKO cells.

Given prior evidence suggesting functional interplay between class I myosins and non-muscle myosin IIA (NM2A) during phagocytosis (Barger et al., 2022), we next examined how loss of Myo1e/f affects NM2 organization within the phagocytic cup. To visualize NM2 localization, WT and dKO macrophages were transfected with a GFP-tagged myosin regulatory light chain (RLC), which binds NM2 heavy chains, and subjected to the same phagocytosis assay described above. Cells were fixed 10 minutes after bead addition and stained for F-actin with phalloidin.

In WT macrophages, NM2 localized as a dense band positioned just behind the leading edge of the F-actin–rich cup during most stages of phagocytosis, shifting only after target enclosure (Fig. 8A,C; Fig.S4D-E). NM2 was also enriched at the outer face of the cup, where it did not contact the bead. In contrast, in Myo1e/f-deficient cells, NM2 distribution was markedly altered: the protein appeared diffusely distributed throughout the F-actin–rich region of the cup along the Z-axis rather than forming a discrete basal band (Fig. 8B-C). Despite this difference in vertical organization, the lateral (X-axis) localization along the outer cup surface remained similar to that of WT cells.

These findings indicate that Myo1e/f are required to maintain the spatial distribution of NM2 within the phagocytic cup (Fig. 8D), suggesting that Myo1e/f help coordinate the balance between contractile and protrusive actin networks necessary for efficient cup progression.

Together, these results reveal that Myo1e and Myo1f play a central role in organizing the cytoskeletal architecture of the phagocytic cup. By localizing to podosomes, actin teeth, and the phagocytic ring, Myo1e/f appear to stabilize the spatial organization of F-actin and constrain NM2 distribution during target uptake. In their absence, loss of podosome integrity, premature formation of the phagocytic ring, and dispersion of NM2 coincide with inefficient or stalled cup progression. These findings suggest that Myo1e/f function as key mechanical and organizational linkers that coordinate protrusive and contractile forces required for successful phagocytosis.

### Loss of Myo1e/f leads to an increase in trogocytosis in RAW 2C4.7 cells

Because loss of Myo1e/f delays phagocytic cup closure and promotes premature cup constriction during early engulfment, we hypothesized that these altered cup dynamics may increase pinching off of small portions of target cells, a process known as trogocytosis (Bettadapur et al., 2020). We have previously observed repeated attempts of phagocytes to constrict and ingest small pieces of deformable targets during live-cell imaging, however, due to the self-healing nature of DAAMPs, these attempts were not successful. To directly test the trogocytic activity of RAW macrophages, we used an assay in which fluorescent antibody-coated HL-60 cells were presented to Cell Tracker-labeled macrophages (Fig.9). Both WT and dKO RAW cells formed phagocytic cups and attempted to engulf opsonized HL-60 cells. Cells were fixed 30 min after target addition, revealing small AlexaFluor-488-positive HL-60 fragments within macrophages (Fig.9A,B). To quantify trogocytosis, we measured AlexaFluor-488 fluorescence intensity in intracellular HL-60 fragments localized within CellTracker-labeled macrophages, as described in Materials and Methods. Both the maximum fluorescence intensity and mean intensity of ingested fragments were higher in dKO macrophages than in WT cells, indicating that dKO cells may remove more material per trogocytic event (Fig.9C,D) while the number of fragments per cell was not significantly different between WT and dKO cells (Fig.9E). These findings suggest Myo1e/f may suppress excessive cup constriction and stabilize the cup-target adhesion, thereby promoting complete engulfment over partial target removal (Fig.9F).

**Figure 9.**
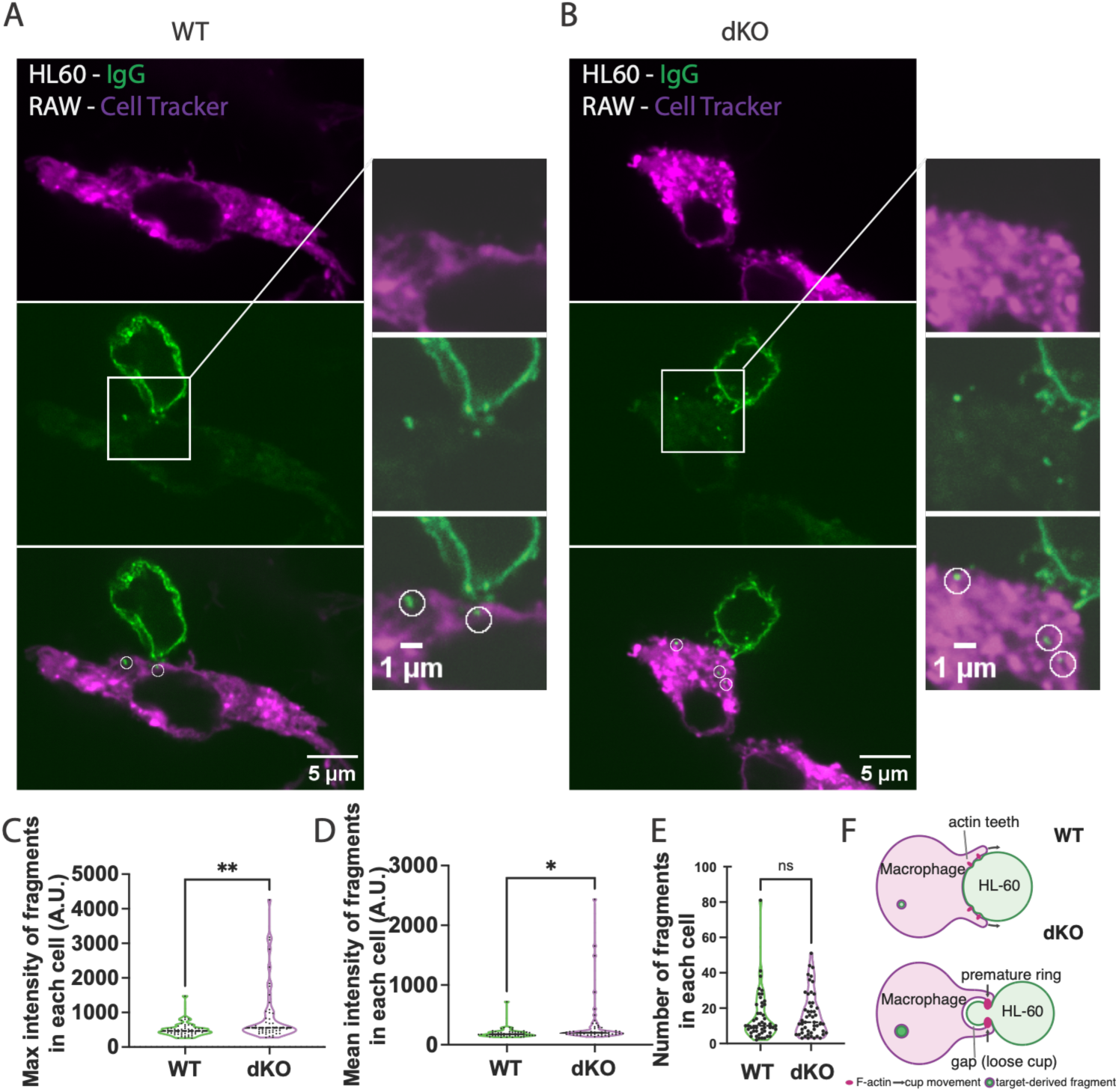
Loss of Myo1e/f increases trogocytosis during antibody-mediated phagocytosis of undifferentiated HL-C0 cells by RAW 2C4.7 macrophages. (A) Representative single confocal section from a 3D image stack of a CellTracker-labeled WT RAW 2C4.7 macrophage (magenta) in contact with an Alexa Fluor 488 anti-biotin-labeled HL-C0 cell (green). Boxed region highlights internalized Alexa Fluor 488-positive target fragments within the macrophage cytoplasm. (B) Representative single confocal section from a 3D image stack of a CellTracker-labeled Myo1e/f dKO RAW 2C4.7 macrophage (magenta) in contact with an Alexa Fluor 488 anti-biotin-labeled HL-C0 cell (green). Boxed region highlights multiple internalized target-derived fluorescent fragments within the macrophage. (C, D) Quantification of the fluorescence intensity of individual internalized target fragments within RAW macrophages. Individual fragments were identified as bright spots with fluorescence intensity at least 50% above the cytosolic background in the IgG channel in control cells. 1 μm circular ROIs centered on each fragment were drawn in ImageJ. Maximum fluorescence intensity (C) and mean fluorescence intensity (D) were determined for each ROI and averaged for each cell. Individual data points in C and D represent cells, with n=50 for WT and n=51 for dKO and N=4. Myo1e/f dKO cells exhibited significantly increased fragment fluorescence intensity compared with WT cells. Statistical significance was determined by unpaired t-test; ** P = 0.0021; * P = 0.0231. (E) Quantification of the number of fragments located inside each cell shown in C and D, replicates are n=50 for WT and n=51 for dKO and N=4. (F) Model illustrating how increased premature cup constriction during early stages of engulfment in Myo1e/f-deficient macrophages may promote pinching and internalization of small target fragments, thereby enhancing trogocytosis (initial drawing created with claude.ai and edited in Adobe Illustrator).

## Discussion

In this study we identified changes in the organization of the phagocytic cup that accompany inefficient engulfment of IgG-opsonized targets in Myo1e/f-deficient RAW macrophages. Formation of discrete actin nodes at the base and rim of the phagocytic cup is markedly reduced, whereas continuous actin rings and pronounced target deformation are prominently observed and occur more frequently before half-engulfment. Live imaging further revealed examples of early target constriction followed by cup retraction, with repeated constriction/retraction cycles. These observations suggest that coordination between local protrusive actin structures, actin ring organization, and cup progression is important for productive phagocytosis. We propose that Myo1e/f help maintain this coordination, allowing target deformation to accompany sustained cup advancement.

Our findings indicate that Myo1e/f contribute to phagocytosis through two complementary mechanisms. First, they regulate the spatiotemporal organization of actin within the phagocytic cup, promoting efficient target adhesion and the coordinated formation of protrusive structures such as actin teeth. Second, they help balance the generation of protrusive and contractile forces within the cup, ensuring smooth progression and cup closure. In the absence of Myo1e/f, this balance shifts from the formation and maintenance of individual podosome-like structures to premature assembly of actin rings, resulting in stalled or aborted phagocytic events, likely due to excessive contractile forces exerted prior to 50% engulfment. Together, these observations suggest that Myo1e/f act as key organizers of cytoskeletal transitions during target engulfment.

Our results extend our previous observations that identified Myo1e/f-positive actin teeth and myosin-II–dependent constriction as key force-generating mechanisms during phagocytosis and implicated Myo1e/f in phagocytic cup closure (Barger et al., 2019; Barger et al., 2022). Using microparticle traction force microscopy and WT RAW cells we have previously demonstrated that actin teeth, enriched in Myo1e/f, exert normal/protrusive forces on the target surface, and that target deformation by protrusive forces is reduced following Arp2/3 inhibition (Vorselen et al., 2021). Primary mouse BMDMs also exhibit protrusive forces leading to phagocytic target deformation, however, the baseline cup architecture differs between macrophage systems: in WT RAW cells, F-actin is concentrated predominantly at the cup rim in the form of discrete teeth (Barger et al., 2019), whereas in WT BMDMs it is distributed more broadly throughout the cup (Barger et al., 2022).

Importantly, earlier work (Ostrowski et al., 2019) showed that during frustrated (planar) phagocytosis the region of closest apposition to the IgG-coated surface coincides with a ring of podosome-like structures rather than the leading edge of the cell and proposed that these actin-rich structures help maintain close membrane-target contact. Consistent with and extending this idea, Myo1e/f-deficient RAW cells examined in the current study were less likely to form podosome-like actin structures, yet retained the ability to deform beads through constriction of the phagocytic rim. Together, these findings suggest that the principal role of actin teeth may be less in bulk target constriction than previously emphasized, and more in maintaining close membrane-target apposition and stabilizing productive protrusive adhesions during cup extension. Loss of Myo1e/f may impair organization of these structures, biasing the cup toward merging of individual podosome-like structures into rosettes and premature constriction.

This interpretation is supported by recent work from Sopelniak et al (Sopelniak et al., 2026), which showed that disrupting podosome assembly via inhibition of Arp2/3 or Src reduced phagocyte-target attachment and impaired overall uptake efficiency and phagosome maturation during uptake of yeast *Candida auris*. Together, these studies highlight a conserved role for podosome-like protrusions in maintaining mechanical coupling between the macrophage and its target. Our data further identify Myo1e/f as key regulators of this coupling, coordinating podosome formation and timing to sustain adhesion and promote efficient cup progression.

Once actin-rich phagocytic rings were formed, we observed that Myo1e/f localized throughout the inner surface of the phagocytic ring, suggesting that this structure may represent a podosome-derived superstructure similar to the podosome rosettes found on ventral cell surfaces during cell migration, matrix degradation, and frustrated phagocytosis (Cougoule et al., 2005; Cougoule et al., 2010; Herron et al., 2022; Ouderkirk and Krendel, 2014; Pan et al., 2011). Notably, these superstructures often do not contain well-defined individual podosomes, instead, they form through coalescence of podosomes into larger, more continuous actin rosettes and belts. Because Myo1e/f are well-established markers of podosomes, their enrichment supports the idea that the ring may be composed of interconnected podosome-like units adapted for the three-dimensional geometry of phagocytic cups, although we did not directly determine whether this is the case. Unlike the ventral podosome rosettes (Ouderkirk and Krendel, 2014), in which Myo1e/f localize to the outer edge of the adhesion, in phagocytic podosome rings these proteins occupy the inner, bead-facing surface, an inversion consistent with the reversed membrane curvature during target engulfment. While in ventral rosettes these myosins face the cell-matrix interface, in the phagocytic cup they are oriented toward the target-facing invagination, effectively ‘pinning’ the membrane to the bead from the inside out.

Our findings also suggest that phagocytic ring formation and contractile constriction are not, by themselves, sufficient to maintain continuous close membrane–target apposition. Although the phagocytic ring can generate substantial circumferential forces and deform the target, such forces may operate at the mesoscale, effectively tightening the purse-string around the target, but without the local anchoring provided by actin teeth. Without the local anchors, the membrane may fail to follow the precise topography of the target, particularly if increased cortical tension during constriction tends to pull the membrane taut. Thus, actin teeth/podosome-like structures may provide a distinct adhesive-protrusive function, acting as discrete sites that maintain intimate membrane-target contact, sustain receptor engagement, and stabilize productive cup advancement. This interpretation is consistent with prior observations in frustrated phagocytosis (Barger et al., 2019). Myo1e/f loss may uncouple ring constriction from local adhesive apposition, allowing the cup to retain rim-dominant contractile deformation while losing the continuous target contact normally supported by actin teeth.

In WT macrophages, NM2 formed a condensed band just below the phagocytic ring, consistent with the presence of linear actin filaments interconnecting the podosome units, as described in frustrated phagocytosis models (Ostrowski et al., 2019). The spatial organization of Myo1e/f and NM2 within this region indicates that the phagocytic ring can integrate both protrusive and contractile functions. Such dual-force generation may explain why phagocytosis can proceed, albeit less efficiently (Araki et al., 2003; Barger et al., 2019), when either Myo1e/f or NM2 activity is impaired, as each system may partially compensate for the other to drive target constriction and closure.

The diffuse redistribution of NM2 observed in Myo1e/f-deficient macrophages provides a potential explanation for the stalled cup progression captured in our LLSM imaging. Van Zon et al (van Zon et al., 2009) proposed a mathematical model in which phagocytic cup extension halts when the maximum force generated by the actin network fails to exceed the force required to advance the cup around the widest point of the target. Our findings are consistent with such a scenario, although our interpretation differs. We propose that, in WT cells, contractile structures assemble after approximately 50% engulfment, where they cooperate with protrusive actin teeth to promote coordinated cup closure. In contrast, in Myo1e/f-deficient cells, contractility appears prematurely and becomes spatially disorganized. Simultaneously, loss of Myo1e/f impairs actin teeth formation or maintenance and reduces protrusive stabilization of the advancing cup. Thus, contractile forces may be sufficient to deform or locally pinch the target membrane yet insufficiently coordinated to drive continued cup advancement around the particle. The combined loss of protrusive support and organized constriction could therefore produce the repeated initiation and retraction cycles characteristic of Myo1e/f-deficient cells while favoring partial target scission events.

Consistent with this model, Myo1e/f-deficient macrophages exhibited slightly increased trogocytosis of opsonized HL-60 targets. Our observations show that both WT and dKO macrophages were actively engaged in trogocytosis, similarly to the observations of Rollins et al. using BMDMs engineered to phagocytose cancer cells (Rollins et al., 2025) and Cornell et al. (Cornell et al., 2025) who examined trogocytosis in several cell lines, including RAW264.7. dKO cells accumulated greater amounts of internalized target material than WT macrophages. We propose that premature, poorly coordinated rim constriction combined with reduced protrusive stabilization favors pinching and scission of small membrane fragments instead of progressive cup extension around the entire target. In this framework, Myo1e/f normally promote productive phagocytosis not only by supporting cup closure, but also by suppressing unproductive fragmentary ingestion events. This finding highlights that trogocytosis may depend not only on the mechanical properties of the target (Cornell et al., 2025) but also the cytoskeletal organization of the macrophage. Given the prevalence of trogocytosis in 3D environments (Rollins et al., 2025) and its role in affecting the efficacy of cancer immunotherapies (Kim et al., 2025; Taylor and Lindorfer, 2015), it is important to determine factors that regulate trogocytosis. Since local formation of protrusive actin structures has also been proposed to drive pinching-off of target fragments (Cornell et al., 2025; Vorselen et al., 2021), and Rac-dependent protrusions increase bite size in intact tissue (Penfield et al., 2026), the specific roles of Myo1e/f in regulating trogocytosis in complex 3D environments will need to be investigated in future studies.

Premature or poorly coordinated formation of contractile structures in Myo1e/f-deficient macrophages may arise from dysregulated NM2 localization or activation caused by loss of Myo1e-dependent signaling and/or altered adhesion architecture. Disruption of small podosome-like adhesion structures could remove local mechanical or biochemical cues that normally constrain NM2 activity to the appropriate stage and region of the cup. Indeed, in our study of conventional podosomes formed on planar surfaces, we have previously found that individual podosome lifetimes were reduced when Myo1e or Myo1f were knocked down in human macrophages, and that cells deficient in Myo1e/f tended to form aggregated clusters of podosome-like structures, which may reflect increased contractility (Cervero et al., 2025). While the specific pathways linking Myo1e and NM2 have not been fully elucidated, Myo1e has previously been shown to interact with LSP1 at sites of phagocytosis, forming an LSP1-Myo1e complex that promotes cup progression (Maxeiner et al., 2015). LSP1 also regulates the distribution and activity of NM2A at podosomes (Cervero et al., 2018). Thus, Myo1e-LSP1 interaction provides one possible pathway, while loss of podosome-like adhesion itself provides an additional non-mutually exclusive mechanism for NM2 misregulation in the absence of Myo1e/f.

Together, our findings establish Myo1e and Myo1f as central coordinators of cytoskeletal remodeling during FcR-mediated phagocytosis. By regulating the formation and timing of podosome-derived actin structures, these myosins maintain the balance between protrusive and contractile forces required for efficient target engulfment while limiting diversion into partial ingestion pathways such as trogocytosis. A deeper understanding of how unconventional myosins integrate with signaling and adhesion networks will be essential for elucidating the molecular logic of phagocytosis. Future work should focus on identifying binding partners of Myo1e/f and defining how these interactions influence actin dynamics, adhesion turnover, and mechanosensation within the cup. Because phagocytosis is inherently mechanosensitive and Myo1e localization has been shown to respond to membrane tension changes (Smith et al., 2026), exploring how substrate stiffness or target rigidity modulates Myo1e/f function may reveal broader principles governing myosin-dependent force coordination in immune cells. Ultimately, these insights will advance our understanding of how the actin–myosin machinery supports immune surveillance and how its dysregulation contributes to pathologies involving defective phagocytosis.

## Materials and Methods

### Cell Culture

Cells (RAW 264.7 and BMDMs) were cultured in high glucose (4.5g/L) Dulbecco’s Modified Eagle medium (DMEM) (Fisher, MT10013CV) with 10% non-heat inactivated fetal bovine serum (Fisher, FB12999102) and 1% penicillin/streptomycin (Gibco, 152240062) and incubated at 37°C with 5% CO2. Cultures were maintained in tissue culture-treated 100mm dishes. Besides instances where the cells were to be used for transfection, cells were detached from the plates with Trypsin-EDTA 0.25% (Gibco, 25200056).

HL-60 cells were cultured in glutamine-supplemented RPMI-1640 (Cytiva, SH30027.01) with 20% non-heat-inactivated fetal bovine serum (Fisher, FB12999102) and 1% antibiotic-antimycotic (HyClone, SV30079.01) and incubated at 37°C with 5% CO2. Cultures were maintained in tissue culture-treated 100mm dishes below 1x10^6^ cells per mL.

### Western blotting

Cell lysates were generated using cold immunoprecipitation (IP) lysis buffer (50 mM Tris/HCl, pH 7.5; 150 mM NaCl; 1 mM EDTA; 1% Triton X-100 (w/v)) with added protease inhibitor (Thermo Fisher, PI88266) and phosphatase inhibitor (Roche, 04906837001). Cells were scraped into the lysis buffer, mixed with a 5X SDS-PAGE sample buffer (50% glycerol, 25% β-mercaptoethanol, 15% SDS, 0.025% bromophenol blue, 250 mM Tris–HCl, pH 6.8), and boiled for 5 min. Samples were separated on an SDS-PAGE gradient gel (7.5%–15% polyacrylamide) and transferred to PVDF membrane (EMD Millipore, IPVH00010). Total protein staining was performed using SYPRO Ruby blot stain (ThermoFisher) according to manufacturer’s instructions. Membrane was blocked with 5% BSA in TBST (20 mM Tris–HCl, pH 7.5, 150 mM NaCl, 0.1% Tween 20) and the membrane was incubated with primary antibodies. After incubating the membrane with HRP-tagged secondary antibodies, blot was developed using ECL reagent (BioRad, 1705060). The ECL signal was visualized using a Chemidoc MP imaging system (Bio-Rad) and Image Lab analysis software (Bio-Rad). The following primary antibodies were used: homemade rabbit anti-Myo1e (RRID AB_2909514, Fig. S1A), rabbit anti-Myo1e (Sigma-Aldrich, HPA023886, Fig. S1D), mouse anti-Myo1f (Santa-Cruz, sc-376534).

### Macrophage Preparation

All experiments involving mouse bone marrow isolation were performed according to the animal protocol approved by the Upstate Medical University IACUC. Myo1e and Myo1f KO mice were maintained on the C57Bl/6 background and all experiments used age- and sex-matched bone marrow preparations. BMDM isolation and differentiation have been previously described (Barger et al., 2022).

Briefly, bone marrow cells were thawed from stocks stored in liquid nitrogen that were frozen after the initial extraction from mouse femurs. They were then maintained in DMEM with 10% FBS and 20 ng/mL M-CSF at 37°C. The medium was replaced with fresh M-CSF-containing medium at 3 days post thaw and medium lacking M-CSF at 7 days post thaw.

RAW264.7 macrophages (ATCC TIB-71) were thawed from liquid nitrogen stock. They were re-passaged 3 days post thaw before use in any assays. WT cells used in this manuscript range from passage 7 to passage 10, while Myo1e/f dKO cells range from passage 18 to passage 23. Myo1e/f dKO cells were generated by CRISPR-mediated gene editing (Fig.1) by Synthego and subjected to limiting-dilution cloning.

### Transfection

Plasmid DNA was transfected into RAW264.7 cells using the Neon electroporation system (ThermoFisher, MPK10096). Prior to transfection, cells were detached using TrypLE Express (ThermoFisher, 12604021) and brought to a concentration of 4 x 10^5^ cells per experimental well. Cells were then centrifuged (300 x g, 5 min) and resuspended in 100 μL of cold R-buffer (part of MPK10096 kit) per experimental well. Plasmid DNA was then added at a concentration of 15 μg per experimental well. Cell/DNA solutions were then electroporated with the following settings: one 1720V pulse for 20 ms. They were then plated onto an acid washed, 18 mm, glass coverslip in the appropriate well of a 12-well plate with 1 mL of DMEM with 10% heat inactivated FBS (RCD Systems, S11150) without antibiotic/antimycotic. Plates were incubated at 37°C for 24 h prior to use.

### Bead Functionalization

9 μm carboxylate-modified polystyrene beads (Bangs lab, PC07001) and deformable acrylamide-co-acrylic acid microparticles (DAAMPs, 8.8 μm in diameter and ∼1 kPa in stiffness) (Vorselen et al., 2020) were coated with cadaverine-conjugated TetramethylRhodamine (TRITC) (Life technologies, A1318) or AlexaFluor-647 (Thermo Fisher, A30676) and Bovine Serum Albumin (BSA) (Sigma-Aldrich, A3059) for use in phagocytosis assays. Beads were functionalized as in Vorselen et al 2020 (Vorselen et al., 2020). Briefly, beads were diluted to 5% (v/v) concentration and washed twice in activation buffer (100mM MES, 200mM NaCl, pH 6.0) using resuspension and centrifugation (11,000 x g, 5min). They were then mixed at room temperature for 15 minutes in activation buffer supplemented with 40mg/mL 1-ethyl-3-(3-dimethylaminopropyl) carbodiimide, 20 mg/mL N-hydroxysuccinimide (NHS), and 0.1% (v/v) Tween 20. Following activation, beads were pelleted and washed three times in PBS pH 8.0 (adjusted with NaOH) with 0.1% Tween 20. Immediately following the final wash, the beads were resuspended in PBS pH 8.0 containing 20 mg/mL BSA and rotated for 1hr at room temperature. Then cadaverine conjugate was added to a final concentration of 0.2 mM and beads were rotated for an additional hour. Unreacted NHS groups were blocked with 100mM TRIS and 100mM ethanolamine (pH9.0) for 30 minutes with mixing. The beads were then washed three times in PBS, pH 7.4 without Tween 20 and resuspended in the same buffer for storage at 4°C. The day before use, beads were washed twice in sterile PBS and opsonized with 3mg/mL rabbit anti-BSA antibody (MP Biomedicals, 651111) while rotating at 4°C overnight. Beads were then washed three times in PBS (with centrifugation at 3000 x g, 5 min) and resuspended in DMEM without serum or antibiotics.

### Phagocytosis Assay

Cells were plated on 18mm acid-washed glass coverslips in 12-well plates at a density of 2.5 * 10^5^cells/well one day before the experiment. For experiments involving transfection, cells were electroporated at the time of plating using the Neon electroporation system (see Transfection section for details). On the day of the experiment, IgG-coated beads were washed three times with PBS and resuspended in DMEM at a concentration of 2 beads/cell in a final volume of 1 mL per well. The medium in each well was then replaced with the bead suspension. In phagocytic efficiency assays utilizing Halo-tagged constructs, cells were suspended in DMEM with added serum and 0.25 μM JF646 Halo-ligand (Grimm et al., 2015) for one hour prior to bead addition. To synchronize phagocytosis across wells, plates were centrifuged at room temperature for 3 minutes at 300 x g to sediment beads onto the cell surface. Plates were then incubated at 37°C for indicated times. Following incubation, cells were fixed by adding 1 mL of 8% paraformaldehyde (PFA) solution into each well (final concentration 4%) for 10 minutes. Wells were then washed three times with PBS for 5 minutes each.

### Phagocytic Efficiency

Following the procedure outlined in the Phagocytosis Assay section, DyLight-488-conjugated goat-anti-rabbit secondary antibody (Fisher, P35552) (1:1000 dilution in PBS) was added to each well for 1 hour. After the antibody incubation, wells were washed three times in PBS for 5 minutes each. The coverslips were then mounted onto glass microscope slides using ProLong Gold (Thermo-Fisher, P36930) or ProLong Glass (Thermo-Fisher, P36980) anti-fade mounting media.

### Fixed-Cell Phagocytic Cup Staining

Following the procedure outlined in the Phagocytosis Assay section, Alexa-405-conjugated goat-anti-rabbit secondary antibody (Thermo-Fisher, A31556) (1:750 dilution in PBS) was added to each well for 1 hour to label exposed bead surface. Following antibody incubation, the wells were washed 3x in PBS for 5 minutes each. Then Triton X-100 0.25% (v/v) in PBS was added to each well for 3 minutes for permeabilization. The Triton X-100 solution was removed, and 1mL of Alexa-568 Phalloidin (Invitrogen, A12380) (1:1000 dilution in PBS) was added to each well for 1 hour. This was followed by washing with PBS three times for 5 minutes each. The coverslips were then mounted onto glass microscope slides using ProLong Gold or ProLong Glass anti-fade mounting media.

### Trogocytosis Assay

Cells were plated on 18 mm glass coverslips in 12-well plates one day prior to the experiment and maintained in complete growth medium. On the day of the assay, RAW 264.7 macrophages were washed three times with PBS and incubated in DMEM containing 24 μM CellTracker CM-DiI for 90 min to label macrophages. Cells were then washed three times with PBS, replenished with complete medium, and returned to 37°C with 5% CO₂ until use.

HL-60 target cells were collected by centrifugation at 200 × g for 3 min, washed in PBS, and resuspended in PBS containing 0.1 M sodium bicarbonate. For biotinylation, 4 × 10^6^ cells were incubated with 6 μL of 292 mM NHS-biotin for 30 min at room temperature. Cells were then washed three times by centrifugation (200 × g, 3 min) and resuspended in PBS. Biotinylated HL-60 cells were labeled by incubation with Alexa Fluor 488 anti-biotin antibody (40 μL of 200 μg/mL stock) for 1 h at 4°C, followed by three additional washes in PBS.

For trogocytosis assays, 1 × 10^6^ labeled HL-60 cells were added to each coverslip containing CellTracker-labeled RAW macrophages. Plates were centrifuged at 200 × g for 3 min to synchronize target-cell contact and uptake, then incubated at 37°C with 5% CO₂ for 30 min. Following incubation, cells were washed three times with PBS and fixed in 4% paraformaldehyde for 15 min at room temperature. Coverslips were then mounted onto glass slides for imaging.

For quantification, 3D image stacks were manually analyzed in ImageJ. Only cell pairs consisting of a single RAW macrophage in contact with a single HL-60 cell, where the macrophage had not fully engulfed any targets, were included. Alexa Fluor 488-positive signal localized within the CellTracker-positive macrophage volume and spatially separable from the main HL-60 cell body was scored as a trogocytic fragment. A 1 μm circular region of interest was drawn around each fragment in corresponding z-planes, and maximum and mean fluorescence intensity values were recorded for analysis.

### Lattice Light-Sheet Microscopy

LLSM experiments were performed at the Advanced Imaging Center at the HHMI - Janelia Research Campus. RAW264.7 macrophages were cultured on 5 mm round coverslips (Warner Instruments Cs-5R). Coverslips were then mounted on the LLSM sample holder and submerged in FluoroBrite medium (ThermoFisher, A1896701) supplemented with 10% fetal bovine serum and 1% Antibiotic-Antimycotic. Imaging medium was maintained at 37°C and 5% CO2 using a custom incubation system (Okolab). The design of the LLSM is as previously described (Chen et al., 2014). Imaging was performed using a square lattice pattern (NA max: 0.4; NA min: 0.325), a Special Optics 0.65 NA, 3.75 mm working distance water-dipping excitation objective, and 488 nm and 647 nm excitation laser lines (100 mW base power, 4% - 20% acousto-optic tunable filter transmission). Volumetric images (512 x 1024 pixel field of view (FOV) with 201 z steps) were collected while scanning the sample stage laterally at an angle of 31.8° relative to the optical axis of the detection objective (Nikon CFI Apo LWD 25x– Water dipping, 63x magnification, 1.1 NA, 3 mm working distance) with a step size of 0.4 µm and an exposure time of 10 ms. Stacks were collected every 6.24 seconds until manually stopped for each experiment based on the duration of the particular phagocytic event being observed. Emitted fluorescence was filtered through two notch filters (Semrock NF03-488E-25 and NF03-642E-25) and imaged onto a Hamamatsu ORCA-Flash 4.0 v2 sCMOS detector. After data collection, images were deskewed and deconvolved (10 iterations of Richardson-Lucy deconvolution with an experimentally determined point spread function) using a custom analysis pipeline (Hobson et al., 2024). The final voxel size after processing was 0.104 × 0.104 × 0.211 µm.

### Confocal Microscopy

Confocal images were collected using the Nikon Eclipse Ti2e Spinning Disk Confocal Microscope equipped with a CSU-X1 (Yokogawa) with an array of 50-micron pinholes with a spacing of 250 microns. The microscope was equipped with a filter wheel with the following filters: ET455/50M (Chroma), ET525/36M (Chroma), ET605/52M (Chroma), ET705/72M EM (Chroma). Sample excitation was achieved using a LUNF-XL (Nikon) laser combiner and an iChrome MLE-LFA-NI2 (TOPTICA Photonics), 4 laser line source with 405 nm, 488 nm, 561nm, and 640 nm excitations. Fixed cell phagocytic cup images were acquired using the CFI60 PLAN APOCHROMAT LAMBDA D 100X Oil (Nikon) objective lens with type F immersion oil (Nikon, MXA22168). Images were captured using an Orca FusionBT (Hamamatsu) camera. Z-stacks were acquired with a NIDAQ Piezo Z device at 0.2μm steps. Z-stacks were collected starting at the ventral-most surface of the cell to the dorsal-most surface of the bead. Automatic deconvolution of images based on microscope specific processing was completed immediately after initial acquisition using NIS Elements Advanced Research version 5.42.02.

### DIC and Wide Field (WF) Fluorescence Microscopy

DIC and WF images for the phagocytic efficiency assays were obtained using a Nikon Eclipse TE2000-E2 Microscope system equipped with a Photometric Prime 95B scientific CMOS camera. Images were acquired using a 20x magnification, Plan Apo, WD 1.0, NA 0.75 objective, using NIS-Elements advanced research software version 4.60.00 64-bit.

### Phagocytic Efficiency Analysis

Widefield fluorescence images were used to determine the approximate extent of bead coverage by macrophages using the following ImageJ analysis pipeline. Images of the bead surface (red channel) and secondary IgG labeling (green channel) were smoothed, followed by using MaxEntropy automatic thresholding in ImageJ. Output from these steps generated masks of the overall bead surface and of the secondary IgG-coated areas where beads were exposed to the medium and not in complete contact with a macrophage. The Fill Holes operation was then run on the masked images to fill in any apparent holes. After this step, ImageJ Watershed operation was utilized to separate any masked objects in contact with each other. Once beads were detected, the Analyze Particles tool was used to identify masked beads that were within the expected range of particle size and to generate a list of ROIs corresponding to the beads. Finally, the resulting list of ROIs was then manually curated to include only the beads that were in focus, circular, and in contact with a cell. For each bead, mean pixel values were measured within the same ROI in the processed binary masks of the bead and extracellular secondary-antibody labeling. The ratio of these mean values represents the secondary-antibody-positive mask area relative to the total bead-mask area. Estimated macrophage coverage of beads was calculated from the ratio of the mean pixel value of secondary-antibody-mask value to the bead-mask value eq 1 and used as a two-dimensional proxy for phagocytic cup coverage.

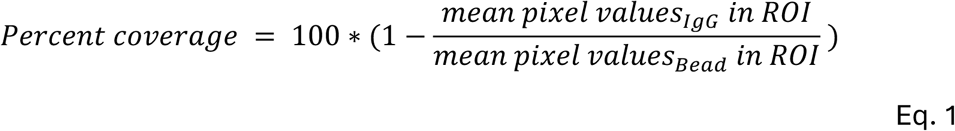

The resulting values produced a bimodal distribution with most bead ROIs being either in the “mostly covered” or “mostly not covered” bins (Fig. S1B). For subsequent numerical analysis, we defined the “fully-eaten” category in the range of 85%-100% coverage and the “in contact with a macrophage but likely not in a phagocytic cup or eaten” in the range of 0%-15% and expressed % phagocytosis as in eq 2.

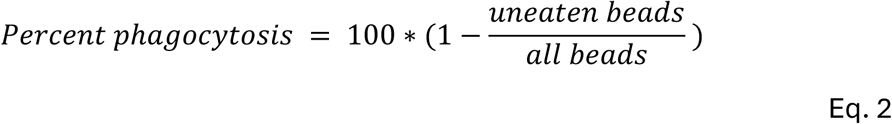

Figures were made using GraphPad Prism software (RRID_SCR_002798). Statistical calculations were performed using built-in analysis operations, with GraphPad Prism 10.

### ChimeraX data visualization

Deconvolved image stacks were visualized in UCSF ChimeraX (Goddard et al., 2018; Meng et al., 2023). From the volumetric intensity data, 3D bead surfaces were rendered as either wire-frame meshes or opaque isosurfaces to observe IgG-coated beads undergoing phagocytosis. Surface curvature was quantified using the ChimeraX *Measure Convexity* function, which reports surface curvature derived from the triangulated surface geometry per vertex. Local curvature values (steradians) were averaged across groups of 10 neighboring vertices to reduce noise and used to generate color-mapped isosurfaces as needed. Additionally, the visualization of adjacent cellular components, such as Lifeact-labeled actin filaments, was performed using maximum intensity projections (MIPs) along the viewing axis.

For preparation of representative time-lapse images and 3D renderings, selected regions of interest were processed in UCSF ChimeraX. The volume region and erase functions were used to restrict displayed volumes to regions containing the cells and DAAMPs of interest. The surface dust function was applied to remove small disconnected voxel clusters arising from background noise in isosurface renderings. These processing steps were used for visualization purposes only and were not applied to raw data used for quantitative analysis. Figures subjected to cropping and/or surface dust filtering include Figs. 3, 5, Supplementary Figs. 2 and 3, and associated supplementary videos.

### DAAM Particle Analysis

Deconvolved images from LLSM and confocal microscopy images were analyzed using custom MATLAB functions adapted from Vorselen et al (Vorselen et al., 2021). Briefly, image stacks of antibody-coated beads were reconstructed in 3D using particle edge coordinates determined with sub-pixel accuracy through the Gaussian fitting of radial intensity profiles as previously described (Vorselen et al., 2021). Prior to reconstruction, image stacks were smoothed to reduce high-frequency noise. The resulting particle surface was resampled with approximately equidistant spacing between neighboring vertices with a separation distance of 250 nm. From the reconstructed triangulation of the particle, we compute its triangulated surface area, S, and volume, V, for subsequent calculation of particle sphericity, fi given by the following equation,

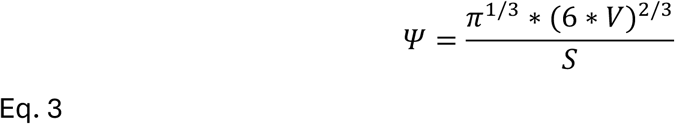

Lastly, the fraction of the particle engulfed is determined through mapping the signal of the fluorescently-stained free (non-engulfed) particle surface onto the reconstructed edge coordinates. To do so, maximum-intensity projections of the smoothed free-surface stain channel (smoothing factor = 0.5), were registered to the triangulated particle’s edge coordinates within a tolerance of 1 µm, followed by masking as described by Vorselen et al (Vorselen et al., 2020).

### Fixed Cup Structural Analysis

Data for Figure 6 A,B,E,F, and G were gathered by manual scoring by an observer blinded to the sample identity using qualitative pre-defined criteria from 3D reconstructed cups visualized in both ImageJ and ChimeraX software. Cells were scored positive for phagocytic rings when a circular band of highly enriched F-actin signal was observed in the phagocytic cup. Basal podosomes were defined as high-intensity F-actin puncta at the bottom of the cup that measured from 0.25-1.0 μm in diameter using a line scan. If a cup had at least one punctum that met the above criteria, it was marked as positive for basal podosomes. Phagocytic teeth were defined as distinct, high-intensity F-actin signals within the body of the phagocytic cup which protruded into the bead-containing region. Loose cups were defined as phagocytic cups which had the presence of secondary antibody signal below the leading edge of F-actin at the rim of the cup. Graphs were made using GraphPad Prism software.

### Schematic drawings

schematic drawing in Fig.9 was initially drafted using claude.ai and manually edited in Adobe Illustrator.

## Acknowledgments

This work was supported by the National Institute of General Medical Sciences of the National Institutes of Health under the Award R01GM138652 to M.K. The Advanced Imaging Center at the Janelia Research Campus is generously supported by the Howard Hughes Medical Institute.

---

**Supplemental Fig. 1.**
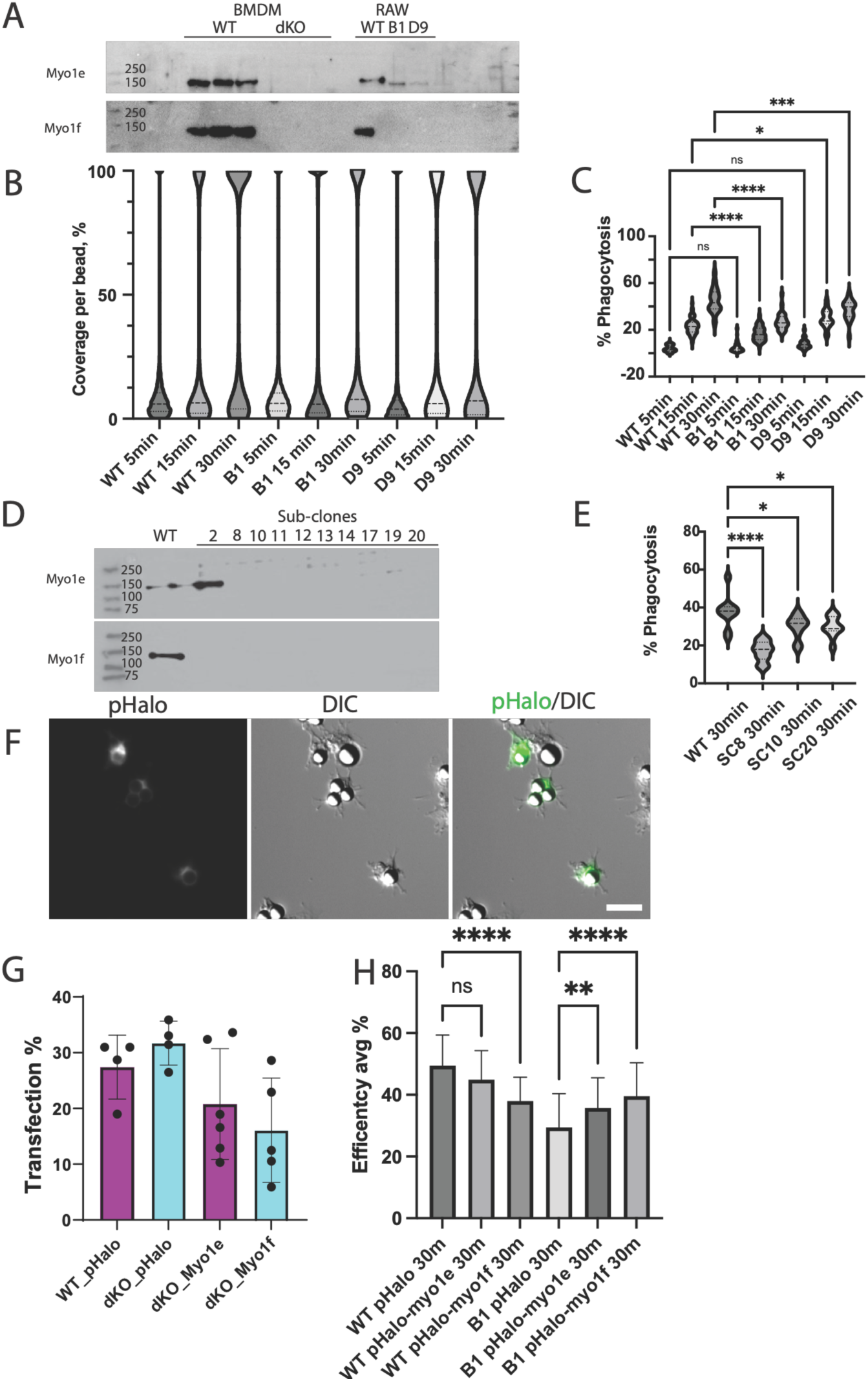
Isolation and testing of additional dKO clones and myosin re-expression experiments. **(A)** Anti-Myo1e or anti-Myo1f Western blot analysis of the two dKO clones provided by Synthego. The Myo1e/f expression in two RAW264.7 dKO clones produced by Synthego (B1 and D9) was compared to RAW264.7 WT as well as BMDM WT and dKO cells. **(B,C)** Time-course of phagocytic uptake of 9 μm polystyrene beads comparing the uptake by WT RAW264.7 and the two Synthego-generated dKO clones B1 and D9. (**B**) Analysis of the distribution of individual beads after secondary antibody labeling that marks the exposed (cell-free) bead surface. % coverage indicates the portion of the bead that is covered by cells and therefore not accessible to secondary antibodies. Replicates graphed of each bead in the dataset n are as follows, WT 5min n= 5684, N=3; WT 15min n= 6595, N=3; WT 30min n= 5707, N=3; B1 5min n= 5928, N=3; B1 15min n= 6944, N=3; B1 30min n= 7461, N=3; D9 5min n= 1156, N=1; D9 15min n= 1042, N=1; D9 30min n= 1011, N=1. (**C**) A violin plot shows the distribution of phagocytic uptake % for multiple fields of view at each of the three indicated time points for each cell line. Both dKO clones B1 and D9 display reduced phagocytic uptake at 15-and 30-minute time points when compared to WT cells. Replicates graphed of each field of view n are as follows, WT 5min n= 58, N=3; WT 15min n= 60, N=3; WT 30min n= 59, N=3; B1 5min n= 54, N=3; B1 15min n= 60, N=3; B1 30min n= 60, N=3; D9 5min n= 20, N=1; D9 15min n= 20, N=1; D9 30min n= 20, N=1. Statistical significance drawn using one-way ANOVA with pairwise comparison tests; * P<0.05, *** P<0.001, **** P< 0.0001. **(D)** Analysis of additional subclones of dKO RAW264.7 macrophages using anti-Myo1e or anti-Myo1f Western blotting. Several monoclonal cell populations, subcloned by dilution from a CRISPR-edited RAW264.7 pool, were examined for the presence of Myo1e or Myo1f using antibodies HPA023886 for Myo1e and sc-376534 for Myo1f. **(E)** A violin plot showing the distribution of phagocytic uptake efficiency values at the 30-min timepoint for multiple fields of view for selected subclones from panel D. Data points correspond to fields of view, n = 10 and N=1. Statistical significance drawn using one-way ANOVA with pairwise comparison tests. * P<0.05, **** P< 0.0001, n=10 FOV, N = 1. **(F)** Montage illustrating transfection of RAW264.7 macrophages with pHalo-tagged constructs for phagocytic efficiency experiments. Scale bar, 25 μm. (**G**) Transfection efficiency observed in a typical re-expression experiment. Cells expressing pHalo-tagged empty-vector or Myo1e/f constructs were identified by comparing the number of cells that had detectable Halo-ligand signal to the total number of cells in the field of view. **(H)** Mean phagocytic uptake per field of view at the 30-min timepoint in WT or dKO (clone B1) macrophages transfected with the empty pHalo vector or Halo-tagged Myo1e or Myo1f. Error bars indicate standard deviation. Re-expression of Myo1e or Myo1f in dKO cells resulted in an increase in phagocytic uptake while expression of Halo-tagged Myo1e or Myo1f in WT cells reduced phagocytic uptake. Replicates graphed of each field of view n are as follows, WT pHalo 30m n = 60, N= 2; WT pHalo-myo1e 30m n = 39, N= 1; WT pHalo-myo1f 30m n = 30, N= 1; dKO pHalo 30m n = 69, N= 2; dKO pHalomyo1e 30m n = 70, N= 2; dKO pHalo-myo1f 30m n = 70, N= 2. Statistical significance drawn using one-way ANOVA with pairwise comparison tests. **P<0.01, **** P<0.0001

**Supplemental Fig. 2. (related to Fig. 3).**
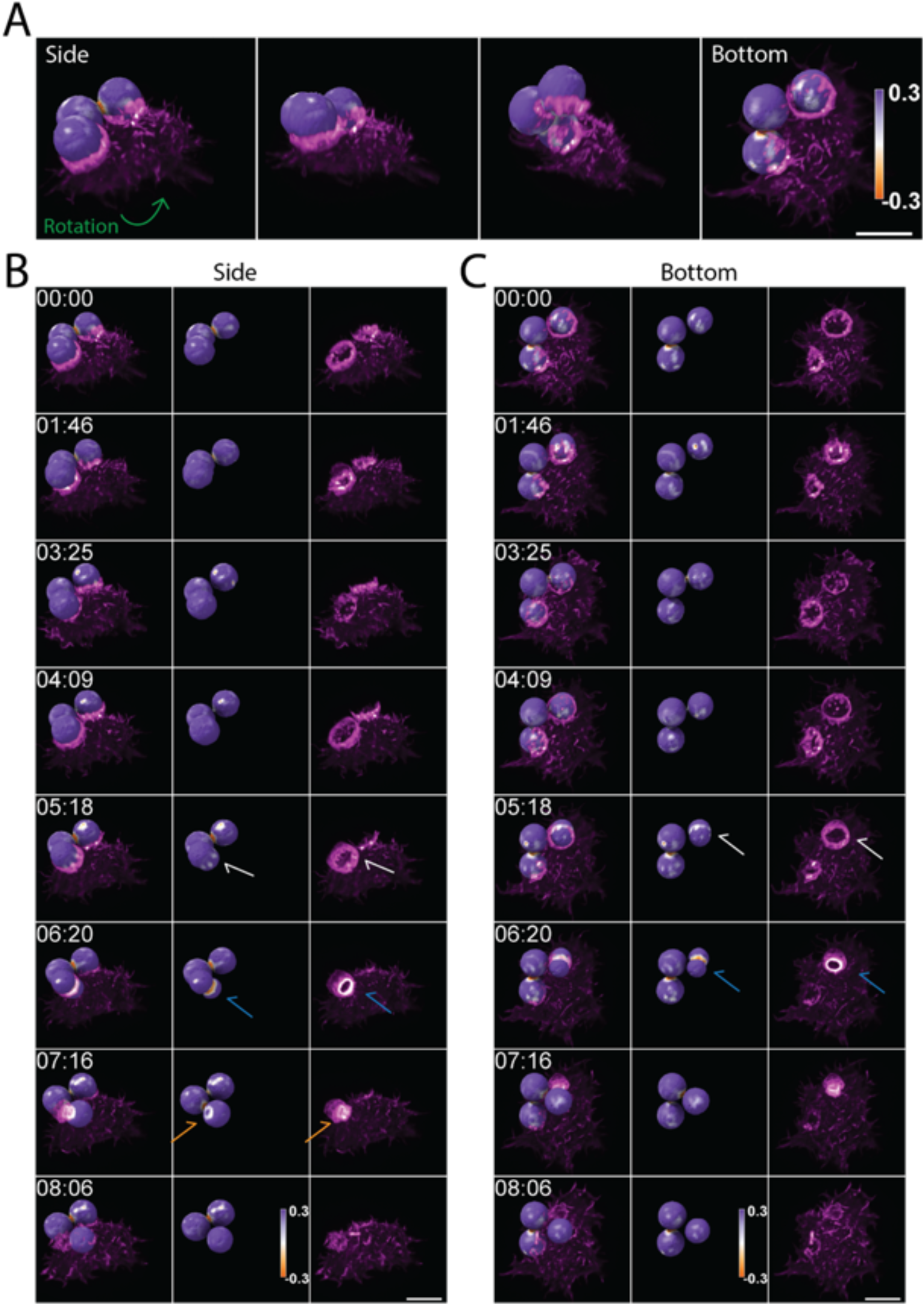
Additional example of actin-driven particle deformation during FcγR mediated phagocytosis. **(A)** The initial frame of an LLSM time-lapse movie showing WT RAW 264.7 macrophages expressing mEmerald-Lifeact engaging several antibody-coated DAAMPs. Lifeact signal is displayed as a magenta-to-white MIP, while DAAMP surfaces are rendered as opaque isosurfaces with a diverging color palette encoding local surface curvature (purple to orange, with white corresponding to zero curvature). The montage shows three progressively rotated side views of the same field of view (left to right), followed by a bottom view. **(B-C)** Individual frames from the same time series show two different viewing angles of the WT macrophages internalizing antibody-coated DAAMPs. During particle uptake, actin dynamics associated with phagosome formation include the appearance of podosome-like structures that induce local indentations in the DAAMP surface (white arrows). These podosome-like structures subsequently coalesce into a ring-shaped structure (blue arrows), which induces deep indentations in the DAAMP. During later stages of internalization, actin pools into a localized spot that continues to indent the particle until phagocytosis is complete (orange arrows). Scale bars, 10μm.

**Supplemental Fig. 3 (related to Fig. 5):**
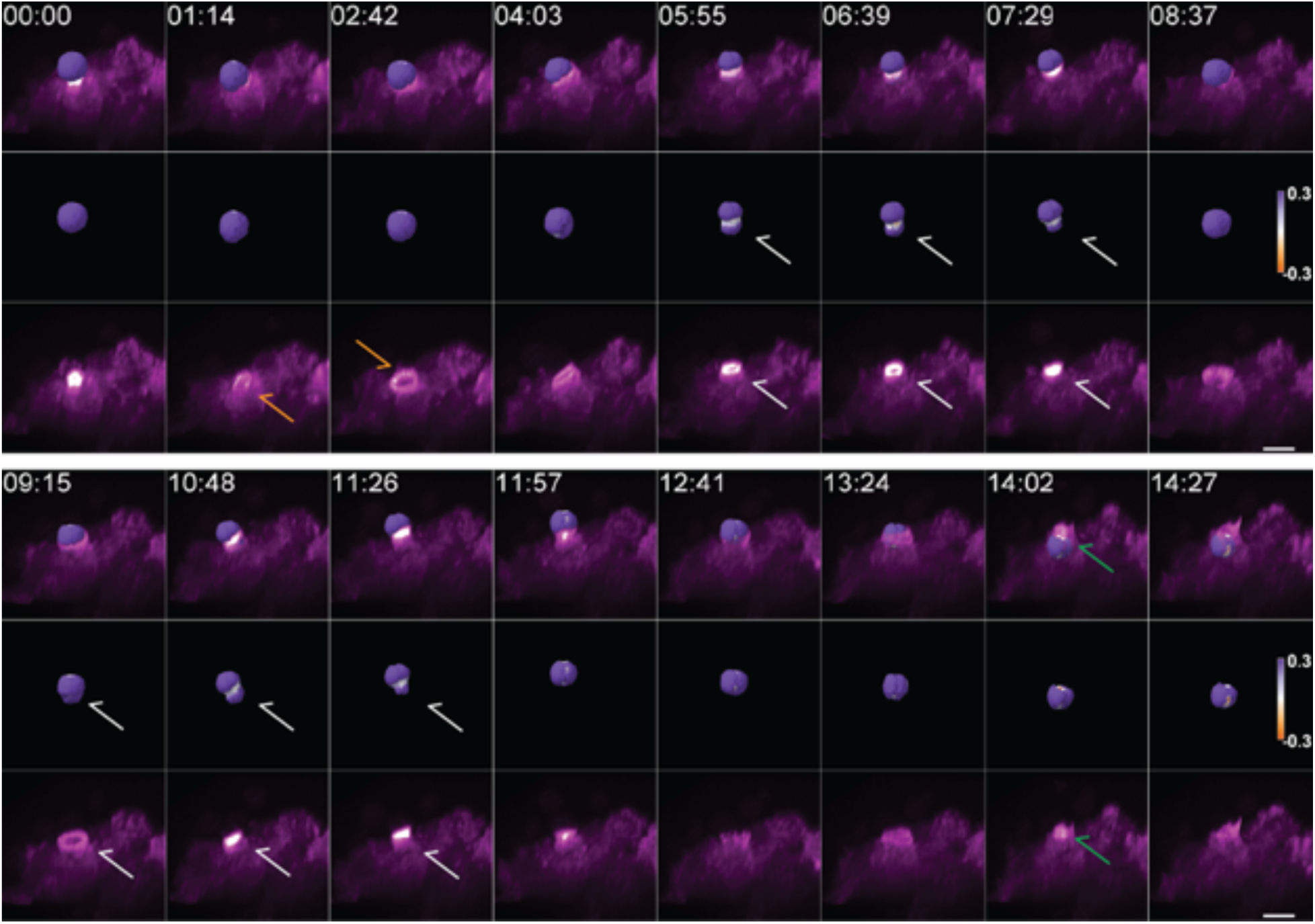
Myo1e/f dKO macrophage repeatedly pinches IgG-coated DAAMPs during FcγR-mediated phagocytic uptake. Time series of a dKO RAW 264.7 macrophage expressing mEmerald–Lifeact, (magenta-white vMIP), attempting to eat an IgG-coated DAAMP rendered as an opaque isosurface with a diverging purple-orange colormap encoding local surface curvature (steradians). Early in the sequence, a localized rosette of F-actin–rich, podosome-like structures forms beneath the DAAMP. This superstructure subsequently disassembles, and a phagocytic cup containing podosome-like puncta becomes evident (orange arrows). Repeated failed uptake attempts are observed when podosome-like structures coalesce into a ring-like configuration that constricts the DAAMP (white arrows) before sliding back and relaxing the constriction. In contrast, successful uptake is later observed during efficient progression and coalescence of F-actin-rich podosome-like structures above the upper hemisphere of the DAAMP (green arrows). Scale bar, 10μm. This time series was cropped to the region of interest and processed with surface dust filtering to display only the bead of interest.

**Supplemental Fig. 4:**
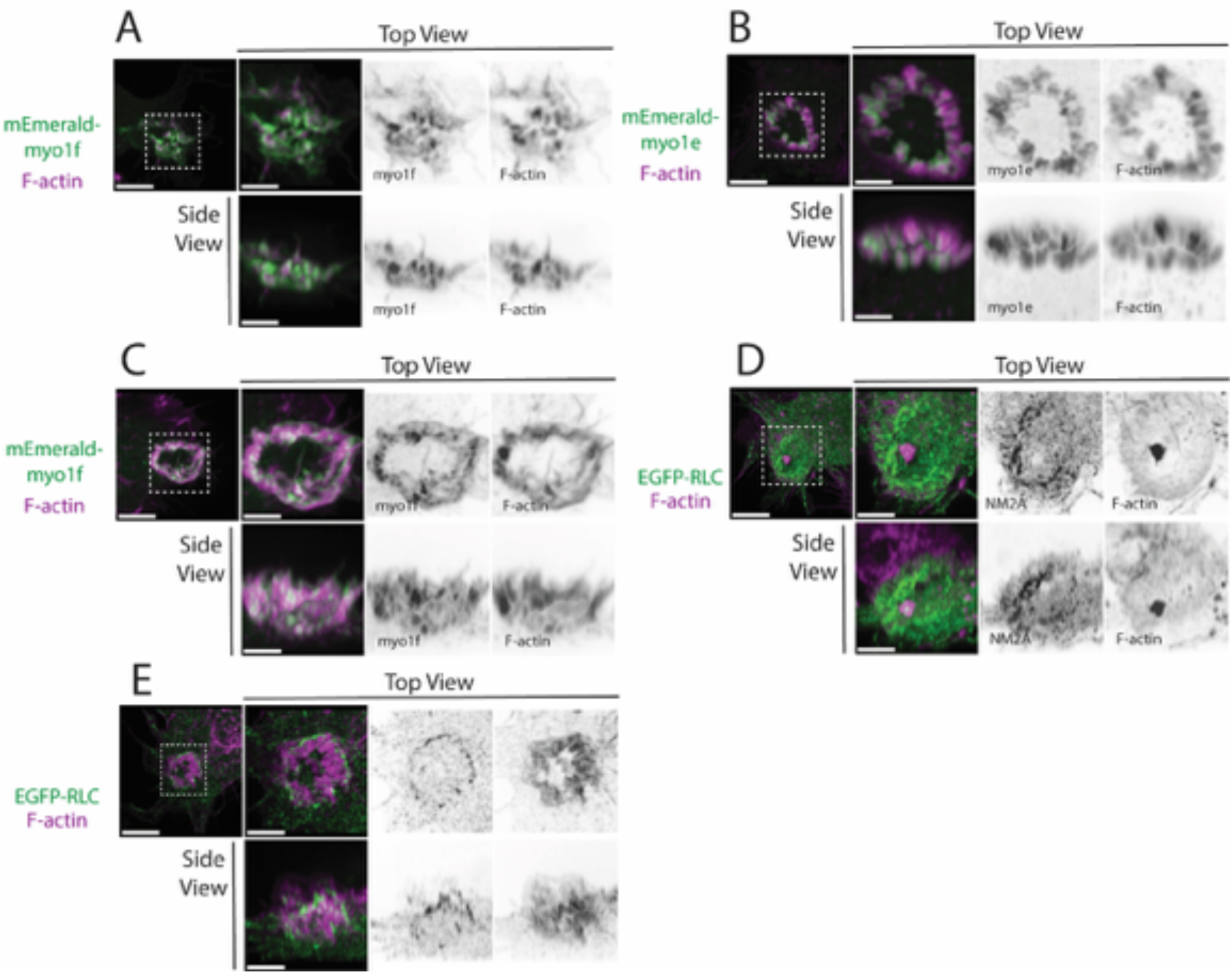
Additional examples of Myo1e, Myo1f, and NM2 localization in phagocytic cups. **(A)** Myo1f localization to basal podosomes during early stages of cup formation. mEmerald-Myo1f (green) localized to the tips of phalloidin-labeled F-actin hotspots (magenta) in basal podosomes. **(B-C)** Myo1e and Myo1f localization to phagocytic actin teeth. mEmerald-Myo1e/f (green) localized to the tips of phalloidin-labeled F-actin (magenta) in actin teeth. **(D)** NM2 localization relative to basal podosomes. EGFP-RLC labeled NM2 (green) did not colocalize with F-actin (magenta) in basal podosomes but was seen throughout the base of the cup. **(E)** NM2 localization relative to phagocytic actin teeth. EGFP-RLC labeled NM2 (green) did not colocalize with F-actin (magenta) in actin teeth but was found in a condensed band at the base of the cup. All panels are RAW264.7 WT macrophages. Scale bars, 5μm, zoom scale bars, 2μm.

**Supplemental Fig. 5 (related to Fig.6):**
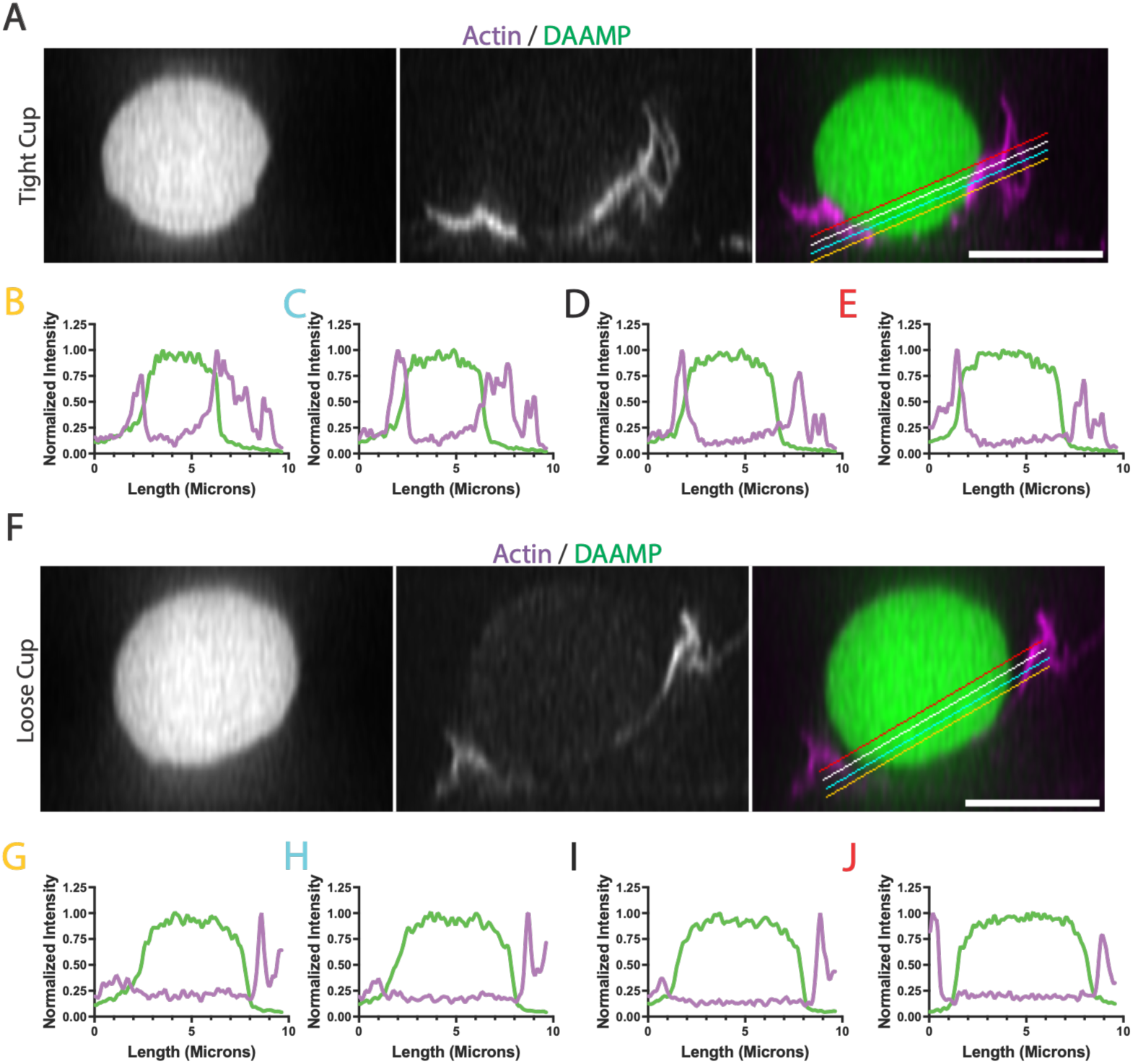
Examples of tight or loose phagocytic cups quantified in Fig.6. **(A, F)** Orthogonal (XZ) view of a phagocytic cup derived from a confocal Z-stack is shown in each montage. Actin is shown in grayscale (middle panel) or magenta (right panel), DAAMP is shown in grayscale (left panel) or green (right panel). Cells in which actin-rich phagocytic cups were in close contact with the bead were scored as “tight cups” (A) while the cups that displayed widely open actin lamellipodia and maintained a gap between the bead surface and cup rim were scored as “loose cups” (F). Scale bars, 10μm. B-E, G-J – line scans showing fluorescence intensity of actin (magenta) or DAAMP (green) along the lines indicated in A and F. Panel labels in B,C,E,G,H,J are color-coded to match the lines in A and F. Line scans in B-E show close apposition between actin signal and the edges of the phagocytic target while graphs in G-J reveal sharp boundaries or gaps between the actin and DAAMP signals.

